# RAGE engagement by SARS-CoV-2 enables monocyte infection and underlies COVID-19 severity

**DOI:** 10.1101/2022.05.22.492693

**Authors:** R. Angioni, M. Bonfanti, N. Caporale, R. Sánchez-Rodríguez, F. Munari, A. Savino, D. Buratto, I. Pagani, N. Bertoldi, C. Zanon, P. Ferrari, E. Ricciardelli, C. Putaggio, S. Ghezzi, F. Elli, L. Rotta, F. Iorio, F. Zonta, A. Cattelan, E. Vicenzi, B. Molon, C.E. Villa, A. Viola, G. Testa

## Abstract

The spread of SARS-CoV-2 has fueled the COVID-19 pandemic with its enduring medical and socioeconomic challenges due to subsequent waves and long-term consequences of great concern. Here we charted the molecular basis of COVID-19 pathogenesis, by analysing patients’ immune response at single-cell resolution across disease course and severity. This approach uncovered cell subpopulation-specific dysregulation in COVID-19 across disease course and severity and identified a severity-associated activation of the receptor for advanced glycation endproduct (RAGE) pathway in monocytes. *In vitro* experiments confirmed that monocytes bind the SARS-CoV-2 S1-RBD via RAGE and that RAGE-Spike interactions drive monocyte infection. Our results demonstrate that RAGE is a novel functional receptor of SARS-CoV-2 contributing to COVID-19 severity.

**One-Sentence Summary:** Monocyte SARS-CoV-2 infection via the receptor for advanced glycation endproduct triggers severe COVID-19.

## Introduction

Since the onset of the Coronavirus disease 2019 (COVID-19) pandemic more than 535 million infections and over 6 million deaths have been reported worldwide (as of June 2022) (https://covid19.who.int/). Despite the striking success of vaccines at mitigating disease severity with the ensuing relaxation of restrictions in many countries, the pandemic remains a major global challenge, due to vaccinations campaigns largely incomplete when not effectively halted worldwide, lack of timely access to effective oral treatments for most patients, the evolution of new variants with increasing immune escape and the emergence of post-COVID sequelae at alarming scales. Thus, despite the remarkable progress achieved thus far, the elucidation of the molecular pathogenesis of the disease, especially in its most severe forms, remains an obviously pressing need (*1*).

COVID-19 is caused by the severe acute respiratory syndrome coronavirus 2 (SARS-CoV-2), an enveloped single positive-strand RNA virus closely related to SARS-CoV and MERS viruses, belonging to the family of the Coronaviridae and genus Betacoronavirus (*2*).

Structurally, SARS-CoV-2 is shaped by four structural proteins; Nucleocapsid protein (N), Membrane protein (M), an envelope protein (E), and Spike protein (S) (*3*). S is a trimeric glycoprotein with two functional subunits (S1 and S2) responsible for host recognition and viral-host cell membrane fusion, respectively. Through the binding domain (RBD) of the S1 subunit, exposed upon S2 subunit proteolytic activation, SARS-CoV-2 binds to Angiotensin-converting enzyme 2 (ACE2) in target cells (*4*, *5*). While ACE2 expression can directly account for SARS-CoV-2 pathogenesis in several target organs (*6*, *7*), accumulating evidence indicates that additional receptors may be at play in mediating widespread indirect damage through disruption of tissue-specific vascular and immune homeostasis (*8*–*10*). Likely as a result of the combination of direct and indirect damage, COVID-19 prognosis and its long term sequelae display remarkable heterogeneity, with host genetics, age, sex and several comorbidities, such as obesity, hypertension, and diabetes identified as risk factors of mortality (*11*–*14*). However, the precise molecular mechanisms linking these conditions to a substantial risk for COVID-19 mortality remain to be elucidated.

In general, severe COVID-19 patients show innate immunity deviations (including hyper-activation) associated with an impaired adaptive immune response. The peculiar delayed innate induction, mainly due to compromised type I IFN myeloid response, allows a higher extent of viral replication (*15*), limiting viral clearance and paradoxically aggravating the immunopathological response (*16*). The resulting inflammation, including the massive release of pro-inflammatory mediators that has been referred to as viral sepsis (*17*), underlies in turn the severe complications and poor outcome in COVID-19 patients (*18*).

While single-cell omics studies have already provided important insights on the molecular pathogenesis of COVID-19 (*19*, *20*), much remains to be elucidated about the molecular mechanisms through which the virus induces immune cell dysregulation across disease course and severity. This is particularly the case for the myeloid compartment, which plays pivotal functions in both protective and detrimental antiviral responses (*21*–*23*). Interestingly, though permissive to SARS-CoV-2 infection, circulating monocytes and macrophages do not express ACE2 and both phagocytosis of infected cells and antibody-mediated entry (*24*) have been proposed as possible mechanisms for infection. Importantly,the high resolution investigation of the immune landscape in a pre-vaccine cohort of COVID-19 patients is relevant also for the clinical management of patients for the still large group of vaccine-hesitant subjects. Elucidating immune mechanisms contributing to the COVID19 severity in these groups might pave the way to novel therapeutic options.

Here, we adopted a longitudinal design to characterise at a single cell multi-omics level the temporal and severity dynamics of COVID-19 immune response. Our approach identified a severity-associated activation of the receptor for advanced glycation endproduct (RAGE) pathway in circulating monocytes. Furthermore, we demonstrated that monocytes bind the SARS-CoV-2 S1-RBD via RAGE and that RAGE-Spike interactions drive monocyte infection. Finally, the longitudinal analysis allowed us to identify drugs potentially useful to revert COVID-19 immune dysregulation during the initial stage of the disease, fostering the development of novel therapeutic strategies.

## Results

### Single-cell multi-omic characterization of the immune compartment of a longitudinal cohort of COVID-19 patients

Peripheral blood mononuclear cells (PBMCs) from a cohort of 20 COVID-19 patients were longitudinally sampled at hospital admission, discharge, and 1 month thereafter (Table S1). To investigate the molecular dynamics of COVID-19 across time and disease severity, we adopted a multi-omics pipeline using a multiwell-based single-cell technology (BD Rhapsody) that includes the analysis of PBMCs’ whole transcriptome and surface proteins (Fig. 1A). After standard pre-processing and filtering of low-quality cells (QC: minimum 800 genes and 30% of mitochondrial reads per cell), we normalised and batch corrected data for 143,428 cells with state-of-the-art tools. Subsequently, we integrated data with Harmony (*25*) (Fig S1A-B) and visualised single cells following dimensionality reduction (via PCA and UMAP) and Leiden clustering to characterise the different immune cell types and subtypes present in our cohort. We then followed complementary approaches, including supervised exploration of marker genes and surface proteins (Fig. S1C, Table S3 and Table S4) to annotate the identity of each cluster, including T cells, B cells, myeloid cells, NK cells and progenitor cells (Fig. 1A) (from now on we refer to “cell family” for the identity of the main immune cell types, and “cell subclusters” for the identity of the subpopulation of cells included in each cell family). Patients were homogeneously distributed across sex and age (Fig. 1A-B).

**Fig. 1.**
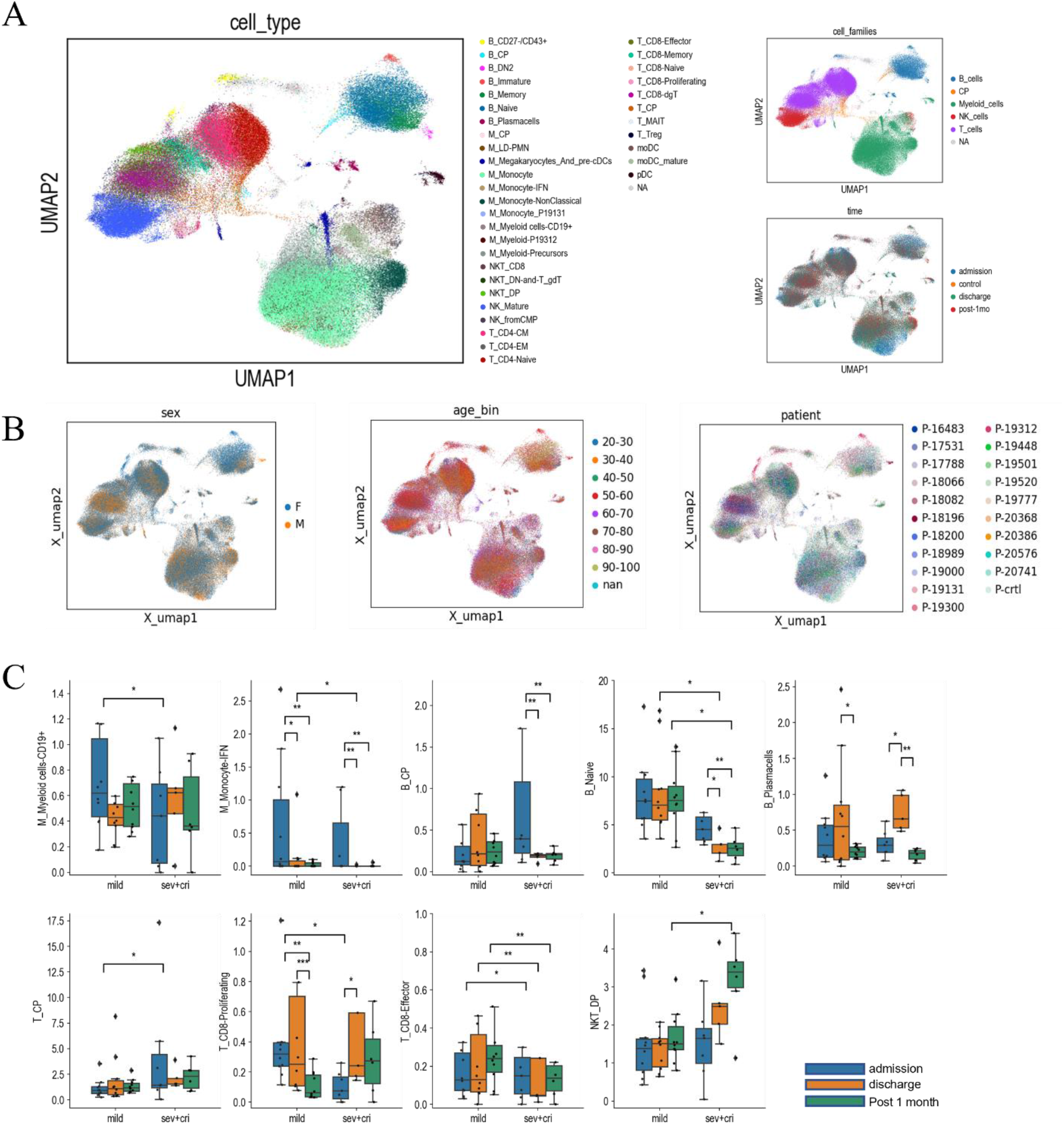
COVID-19 patients single-cell transcriptomic data and cell population enrichment. (**A**) Overview of cell clusters by Uniform manifold approximation projection (UMAP) plots from 143428 single-cell analyses. Main clusters were identified and annotated by family (B_cells, Common Progenitors or CP, Myeloid cells, NK cells, T cells) and then classified in subclusters by specific cell type and their distribution over time (admission, discharge, and post 1 month). (**B**) UMAP plot of the cell coloured by patient sex (left panel), by patient age binned in 10-year intervals (middle panel) and by patient id code (right panel) (**C**) Box-and-whisker plots showing the significant changes of cell type abundance for the COVID-19 samples across time and patient severity. The box shows the dataset quartiles while the whiskers extend to the rest of the distribution, except for “outlier” points that are below/above the first/third quartile with a distance of more than 1.5 times the inter-quartile range. Differential abundance analysis was performed as described in Materials and Methods. *p ≤ 0.05; **p ≤ 0.01: ***p ≤ 0.001.

In order to analyse the impact of SARS-CoV-2 infection on the relative abundance of the different immune cell types and point out differences across disease severity and time, we followed a differential abundance analysis that tests for significant changes in cell composition across conditions (*26*). After quantifying the number of cells both per cell family and per cell subcluster, we identified which clusters were depleted or enriched across disease severity and times (Table S5). Among the cell family clusters, we observed a significant increase of progenitors in mild patients at admission, and a significant decrease of B cells over time, particularly in severe patients (Fig. S2A). When testing for the changes in the subclusters instead, we identified several significant alterations (Fig. 1C). Notably, although the expected reduction of interferon-producing monocytes from admission to later time points was consistent across mild and severe patients, there was an opposite trend of CD8^+^ T effector lymphocytes in mild and severe patients, with an increase in the former and a decrease in the latter. Moreover, the abundance of proliferating CD8^+^ T cells was higher at admission time in mild compared with severe/critical patients. These results highlight the importance of charting the temporal dynamics of the disease at single cell resolution, and are in line with the enrichment of NKT and proliferating CD8^+^ T cells in individuals with more severe infection independently observed by Stephenson et al. (*19*).

To identify the specific longitudinal patterns characterising the gene expression changes induced by SARS-CoV-2 infection in each cell type along the course of the disease, data were further analysed through pseudo-bulk expression profiles, generated by summing counts together for all cells of the same patients and time points, per each cell family cluster, to leverage the statistical rigour of generalised linear models for differential expression analysis (*27*, *28*). Next, we performed a regression analysis along disease time separately for mild and severe patients (Fig. 2A), to define lists of differentially expressed genes (DEGs) (Table S6) showing a specific pattern of expression change over time (admission, discharge, and follow-up), as shown by the heatmaps in Fig S2C. For each cell family and severity, we focused on the DEGs that increased or decreased linearly over time and characterised them through functional enrichment analysis. By systematically comparing the categories that show significant enrichment across all cell families and severities for the linear pattern DEGs, we observed that biological domains related to immune response, cell cycle, metabolism, and translational regulation characterise most of the identified DEGs (Fig. 2B). In detail, we observed significantly enriched pathways in metabolic, mitochondrial, and ribosomal associated gene families in T cells (Fig. 2B, Table S7). Concomitantly, as expected, the functional enrichment analysis revealed a widespread activation of genes belonging to the Immune Response pathway family. Looking more in-depth at the specific differences in the Immune Response pathway of each cell family in severe/critical versus mild patients (Fig. 2C), in line with previous reports (*29*), we observed a strong enrichment in family pathways associated with the IFN signalling and antiviral response, mainly in B, myeloid, and NK cells, with a decreasing trend of expression over time, independently from patients’ severity. On the other hand, in innate cells (Myeloid and NK cell families) the analysis revealed activation of genes involved in the positive or negative immune response regulation in both mild and more severe patients over time. Conversely, in Myeloid and T cells, antigen (Ag) processing and Ag-dependent responses were up-regulated over time, especially in severe patients. Intriguingly, among the significantly enriched pathways, we identified only one category underpinning the direct involvement of a receptor, the GO. 0050786 “RAGE receptor binding” (Fig. 2C).

**Fig. 2.**
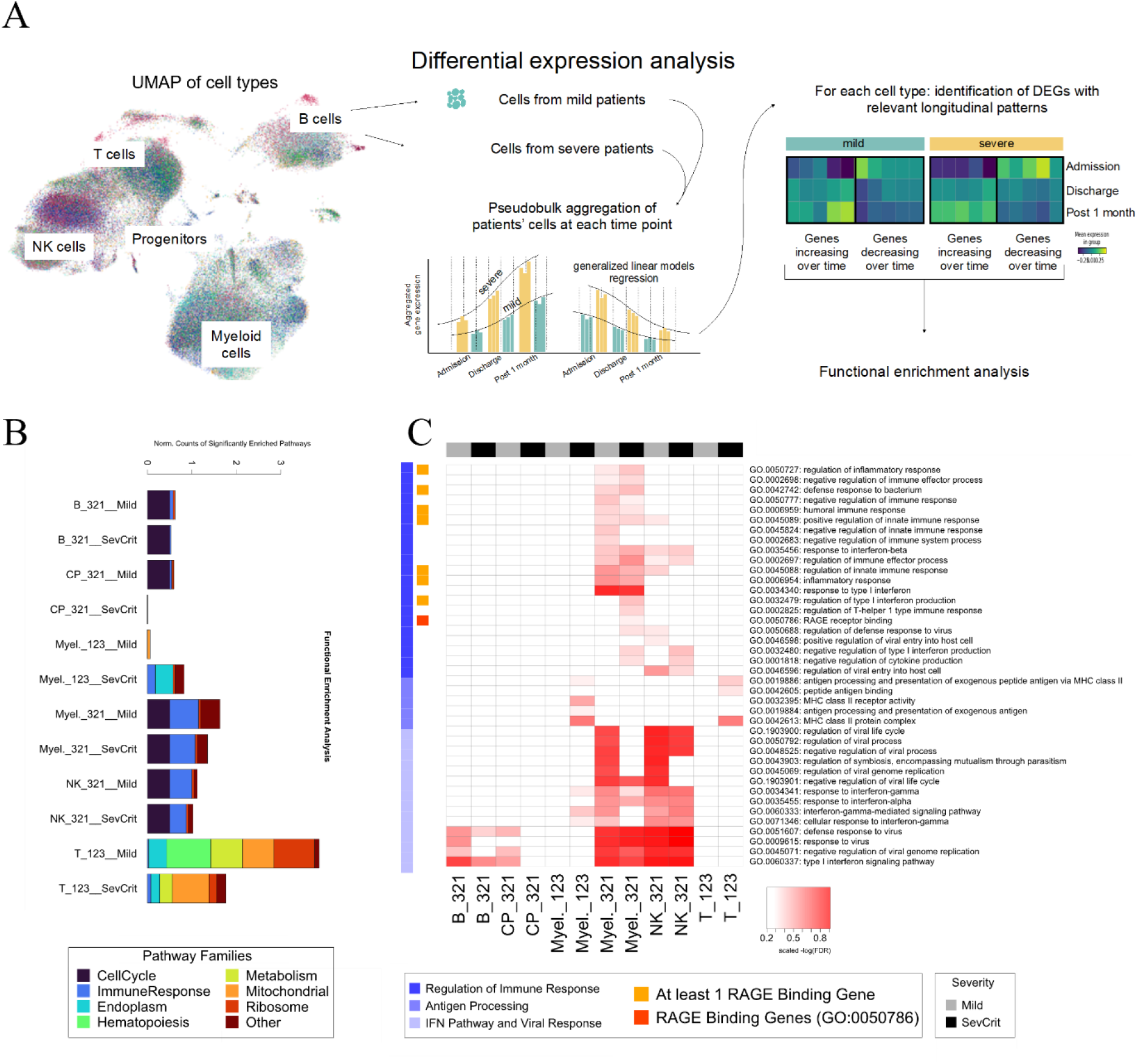
Functional enrichment analysis. (**A**) Schematic representation of the pseudo-bulk differential expression analysis that has been done with the single-cell data. The RNAseq and AbSeq counts for each cell family, patient and sample time-point were summed together to generate pseudo-bulk counts. Separately for the two selected categories of disease severity (mild vs severe+critical), the pseudo-bulk counts were then fitted with a generalised linear model, to identify those genes characterised by a well-defined decreasing or increasing trend of the expression over the sample time-points. These lists of genes were then used as input for the functional enrichment analysis. (**B**) Stacked histograms reporting the counts of significantly enriched pathways (FDR ≤ 0.001 for terms with the number of genes in background between 3 and 500) by cell population, expression trend over time and severity; the counts were normalised by the total number of terms mapped onto each pathway family. Gene sets showing significantly increased or decreased expression trends over time in each cell family were selected in the Mild and SC patients separately. (**C**) Heatmap showing the distribution of the significantly enriched terms related to immunity response among cell population, expression trend over time and severity.

### RAGE pathway activation in Monocytes_IFN at the admission time

RAGE has been implicated in the pathogenesis of several disorders and associated with conditions that predispose to develop severe or critical COVID-19 such as diabetes, hypertension, and ageing (*30*). In our analysis, up-regulation of the RAGE pathway in myeloid cells emerged as a signature of severe/critical patients. Interestingly, myeloid cells are known to be permissive to SARS-CoV entry (*31*) despite the fact that they do not express ACE2, the conventional SARS-CoV and SARS-CoV-2 receptor (*32*, *33*). We thus reasoned that the SARS-CoV-2 might interact with RAGE and trigger signalling events in myeloid cells of COVID-19 patients. We computed the average expression of the genes included in the GO. 0050786 “RAGE receptor binding” category (pathway score (*34*)). We observed that the RAGE pathway score was negligible in all other cell families while much higher in monocytes (Fig. 3A-B), particularly in IFN activated monocytes (Fig. 3B-C). Moreover, we found both a significant decrease of the pathway over our longitudinal timeline and a significant difference in its expression between mild and non-mild patients at admission (Fig. 3D). All single genes of the RAGE pathway plotted in monocytes are depicted in Fig. S3.

**Fig. 3.**
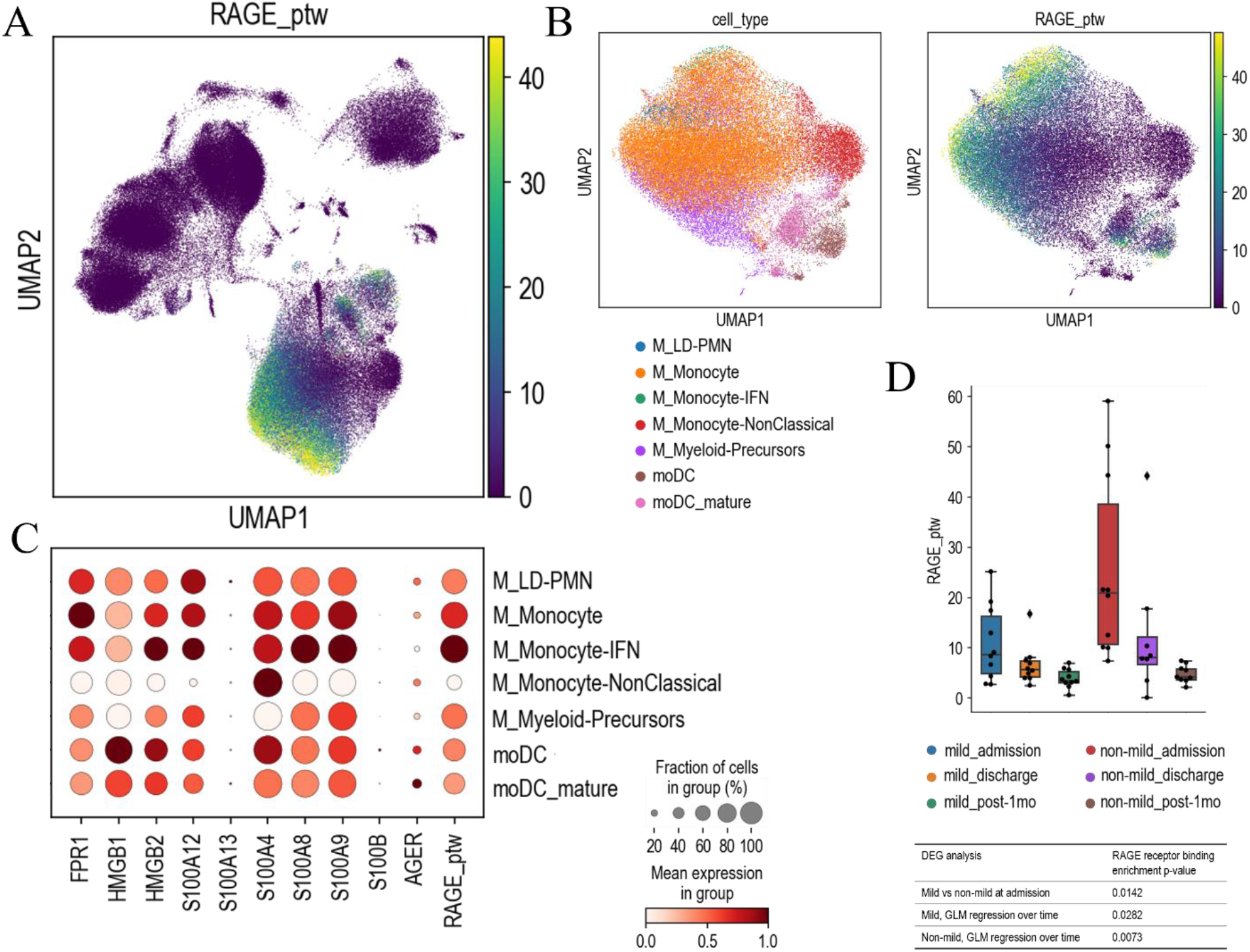
RAGE pathway enrichment. (**A**) UMAP plot of the average expression of the RAGE pathway, defined as the gene score (*34*) of the RAGE receptor binding Gene Ontology Term (GO:0050786). This gene list includes: FPR1, HMGB1, HMGB2, S100A12, S100A13, S100A4, S100A8, S100A9, S100B. (**B**) Close-up view of selected myeloid populations (left panel) with the corresponding RAGE pathway gene score (right panel). (**C**) Dot plot of the average expression values of the GO:0050786 gene list, along with the AGER gene and the RAGE pathway gene score, for each of the selected myeloid populations. Dots are coloured according to the expression value averaged over cells labelled with the same cell type and standardised between 0 and 1 for each variable considered. The size of the dot indicates the fraction of cells within each group with an expression value greater than 0. (**D**) Differential analysis of the RAGE pathway gene score across time and patient severity. The box-and-whisker plot shows the value of the RAGE pathway gene score for all the samples averaged over the cells of the myeloid populations considered. The box and the whiskers are defined analogously to Fig 1B. The table lists some RAGE pathway enrichment p-value, computed as described in Materials and Methods.

Remarkably, we observed that RAGE pathway activation is a reproducible signature validated in all the main publicly available single cell-RNAseq datasets on COVID-19. After selecting blood cells annotated as myeloid from 13 single cell studies previously integrated by Tian et al. in (*20*), we observed that the RAGE receptor binding gene score is consistently different when comparing COVID-19 patients vs healthy controls, and across severity (Fig. S4B). Moreover, we performed pseudo-bulk differential expression analysis and found that DEGs are enriched for the RAGE receptor binding pathway in patients and, among them, for the more severe cases (Fig. S4B). Together, these data uncover RAGE pathway activation as a highly robust signature of SARS-CoV-2 impact on myeloid cells that is associated with COVID-19 severity across cohorts.

### Monocytes bind the SARS-CoV-2 S1-RBD via RAGE

In order to test whether this activation was dependent on the binding of the spike protein with RAGE, potential RAGE and S1-RBD interaction was firstly investigated by generating a computational model of the interaction of S1-RBD to the RAGE receptor (Fig. 4A, Fig. S5A). The model was obtained by molecular docking and further refined using molecular dynamics simulations. Two different models for S1-RBD were obtained using both the original reference sequence first discovered in Wuhan (*35*) and the omicron variant sequence (*36*) that is rapidly spreading worldwide. Docking models were generated using LZerD webserver (*37*), assuming that the interaction is mediated by the RAGE V-domain and by the S1-RBD, as this is the part of the virus that mediates the binding to the ACE2 receptor as well (*38*), and selecting the most probable configuration (highest score). Starting from this configuration we built a full atom model with water and ions (K^+^ and Cl^-^ at 150mM) and ran 5 copies of equilibrium molecular dynamics (MD) simulations for 100ns to confirm the stability of the complex using the protocol described in the Methods. Our results suggest that the binding between the two proteins is possible and stable within the simulated time window and that the interaction is mainly mediated by a salt bridge between S1-RBD Glu471 and RAGE Lys37 and by a network of hydrophobic interactions of S1-RBD residues Leu452, Ile472, Phe490, with RAGE residues Pro33, Val35, Phe62, Leu79, Pro87 (Fig. S5A-B). A visual inspection of the trajectories shows that there are very minor differences between the reference type and the omicron variant (Fig. S5C). Indeed, the only difference in the contact zone for the two variants is represented by the mutation E484A. In the reference type, Glu484 interacts weakly with Arg216 and the backbone of Val35, while in the omicron variant it interacts exclusively with Val35. Computation of the binding free energy confirms that the binding of the S1-RBD to the RAGE receptor for the two variants is indistinguishable within the error range (Fig 4B) and it is much lower than the estimated binding energy between S1-RBD and the human ACE2 receptor, with a ΔΔG = −4.1 Kcal/mol (*39*).

**Fig. 4.**
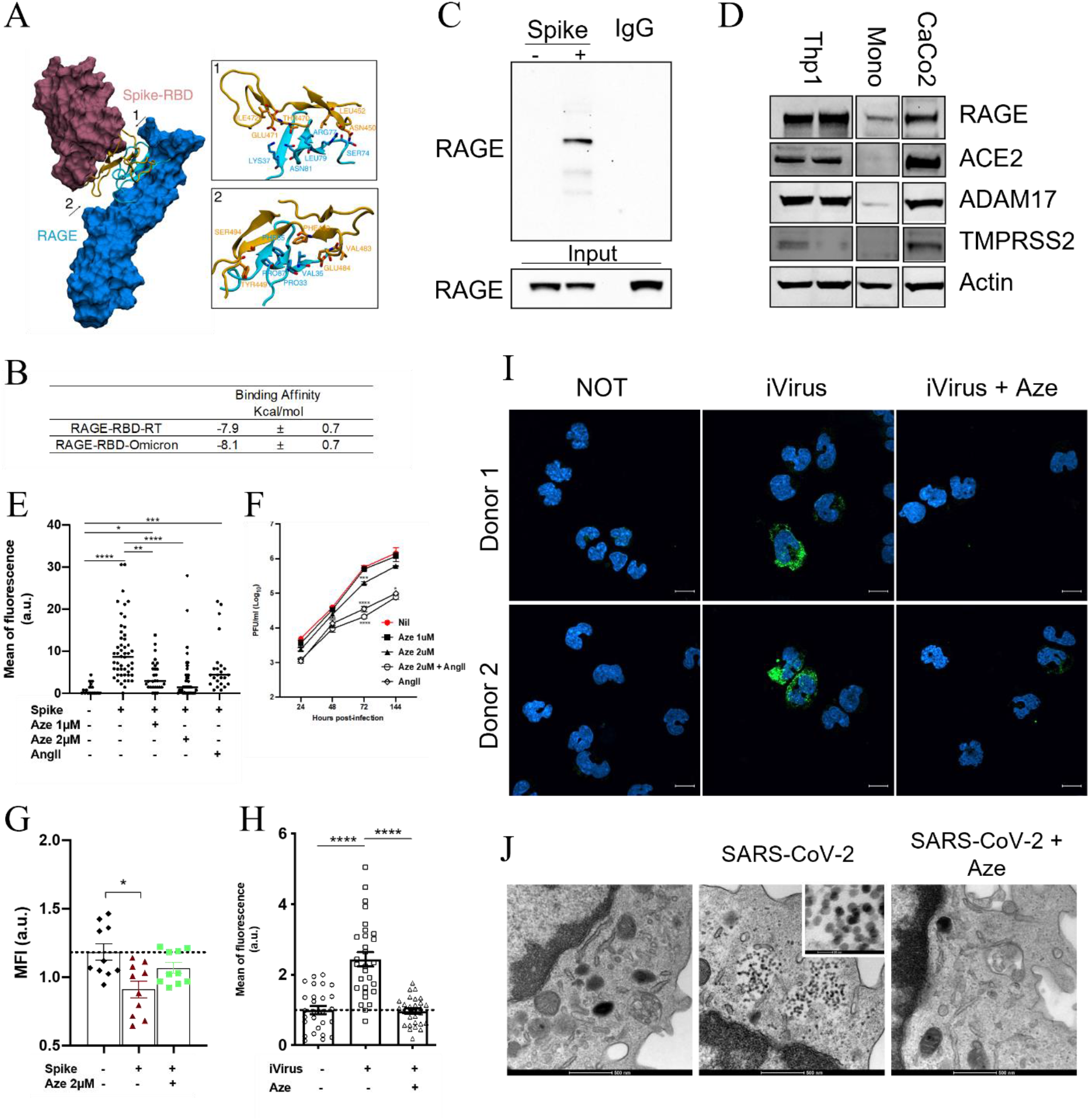
SARS-CoV-2 binds monocytes through an alternative receptor. (**A**) Computational model of the interaction of S1-RBD to the RAGE receptor obtained by molecular docking and further refined using molecular dynamics simulations. Regions of interaction between the two proteins are shown in cartoon representation, and residues important for the binding are explicitly shown from different point of views in sub-panels 1 and 2. (**B**) Computation of the absolute binding free energies of the Reference Type and Omicron Spike RBDs to the RAGE receptor are the same within the errors (see Materials and Methods). (**C**) Co-immunoprecipitation analysis in human peripheral blood monocytes treated with the 100 ng/mL Spike protein. Total proteins were precipitated with an anti-spike antibody and developed with an antibody against RAGE. IgG was used as an antibody specific control. (**D**) Representative Western Blot of RAGE, ACE2, ADAM17 and TMPRSS2 in a monocytic cell line (Thp1), human derived monocytes (Mono) and human carcinoma cell line (CaCo2). Actin was used as a loading control. (**E**) The binding of His-Tag Spike to THP1 cells treated or not with 1-2μM Azeliragon (Aze) was measured as Mean of fluorescence, after 2 hours of stimulation. (**F**) Plaque forming unit (PFU) quantification after THP-1 cells infected with SARS-CoV-2 (0,01 MOI) for 24, 48, 72 and 144 hours in absences or presence of Azeliragon (1 and 2 μM) and Angiotensin II (10μM). (**G**) RAGE internalisation in human-derived monocytes exposed to 100 ng/mL Spike protein alone (+) or upon a pre-treatment with 2μM Azeliragon (+Aze) and measured by flow cytometry after 60 minutes. Data represent the mean fluorescence of RAGE antibody measured by Flow Cytometry. Values were normalised on the RAGE mean of fluorescence of untreated cells (-) at 5 minutes of Spike protein treatment. (**H**) Quantification and (**I**) Representative images of Heat-inactivated SARS-CoV-2 (iVirus) binds human-derived monocytes after 2 hours of incubation. Quantification of fluorescence intensity of Sars-CoV-2 spike protein (green) in monocytes upon infection. Data are presented as mean of fluorescence intensity normalised on Ctr (cells not infected). (**J**) Representative TEM pictures of monocytes treated with replicative SARS-CoV-2 virus (+) MOI 0,1 in the presence or absence of Azeliragon (2μM) pre-treatment. Data are presented as a Mean +/− SEM. Kruskal-Wallis test for multiple comparisons with Dunns post hoc. *p ≤ 0.05; **p ≤ 0.01: ***p ≤ 0.001.

The interaction between SARS-CoV-2 Spike and RAGE predicted by the computational model was verified through co-immunoprecipitation experiments in human peripheral blood monocytes (Fig. 4C). Notably, although the THP1 monocytic cell line express both ACE2 and RAGE, concomitantly with ADAM Metallopeptidase Domain 17 (ADAM17) and Transmembrane Serine Protease 2 (TMPRSS2), primary monocytes do not express ACE2 and TMPRSS2 but express RAGE and ADAM17 proteins (Fig. 4D) (*32*). We analysed the binding of His-Tag Spike to THP1 cells treated with Azeliragon, a small-molecule antagonist of RAGE (*40*), or Angiotensin II, a vasoconstrictor peptide that competes for the ACE2 receptor (Fig. 4E, Fig. S5A). We observed that S1-RBD binding to THP1 cells was hindered by Azeliragon and, as expected, by Angiotensin II treatment. Azeliragon was not toxic to THP1 cells (Fig. S5B) even after 72 hours of treatment. To confirm the RAGE-mediated entry of SARS-CoV-2 in THP1 cells, infections were performed using replicative SARS-CoV-2. significant reduction of viral titers was obtained 72 hours post-infection in 2μM Azeliragon pre-treated THP1 cells (Fig. 4F, Fig. S5C). As expected, consistently with the expression of ACE2 in THP1 cells, Angiotensin II strongly reduced the viral titers in this cell line (Fig. 4F, Fig. S5C). These results demonstrate that both receptors are involved in THP1 cells permissiveness to SARS-CoV-2 entry. To investigate whether SARS-CoV-2 can bind to RAGE independently of ACE2, we moved to primary monocytes, which are characterised by undetectable levels of ACE2 expression. We confirmed that His-Tag Spike protein binds to monocytes and that Azeliragon significantly blocks this interaction (Fig. S5D). Additionally, SARS-CoV-2 spike protein induced RAGE internalisation in primary monocytes, as detected by flow cytometry 30 minutes upon S1-RBD stimulation (Fig. 4G). To further corroborate these data, we incubated freshly isolated human monocytes with a heat-inactivated SARS-CoV-2 (iVirus, VR-1986HK, ATCC). Immunofluorescence staining revealed the presence of viral particles in monocytes, while viral particles were not found in cells treated with Azeliragon (Fig. 4 H-I). The internalisation of replicative SARS-CoV-2 in monocytes was further confirmed through TEM (Fig. 4J, Fig. S7A). In accordance with previous results, 2μM Azeliragon pre-treatment of monocytes reduced the internalisation of infectious SARS-CoV-2 in monocytes (Fig. 4J, Fig. S7A).

### Transcriptional impact of existing and repurposable compounds on RAGE pathway

To probe the potential of available compounds to mimic and potentially revert the transcriptional dynamics underpinning disease course or severity uncovered above, we interrogated the Connectivity Map (CMap) database (*41*), a comprehensive catalogue comprising gene expression profiles of cell lines treated with a panel of ~5,000 compounds. We first tested whether current COVID-19 therapeutics (Dexamethasone, Baricitinib, Ritonavir) were able to mimic the transcriptional changes occurring from admission to post-1month (Table S2). Among the three, only Baricitinib recurrently showed a significant effect on the interrogated COVID-related transcriptional signature (p-value < 0.05 for B cells, NK cells and Myeloid cells). To assess the significance of this result, we further re-performed this analysis following 10,000 randomizations of the CMap drug labels. This highlighted that the probability of retrieving Baricitinib as a significant hit by random chance, due to the recurrence of this drug label in the CMap, is less than 5%. Interestingly, the dissection of Baricitinib’s impact on the RAGE pathway expression confirmed an effect on these genes, with the raw z-scores indicating that Baricitinib is expected to effect a general repression of the RAGE pathway along the disease course, for both mild and severe patients (Fig. 5A).

**Fig. 5.**
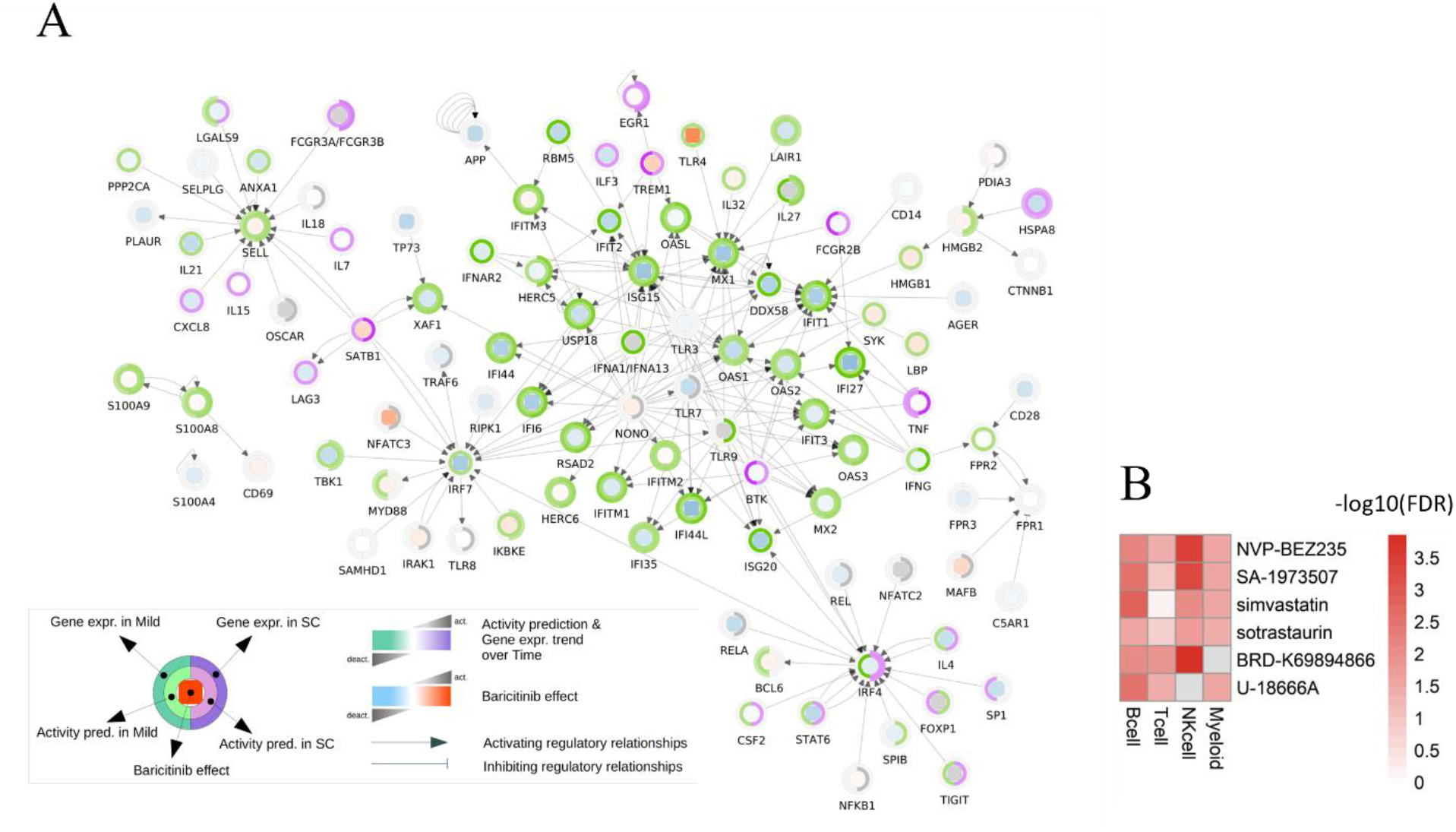
Candidate compounds for COVID-19 treatment based on the pathogenic mechanisms uncovered through our longitudinal study design. (**A**) Extended network of the RAGE receptor binding interactors. In each node three metrics are reported: on the central heatmap the predicted effects of Baricitinib on the gene expression; on the inner circle the predicted overall effect of interactors on the gene expression in Mild (left half-arch) and SC (right half-arch); on the outer circle the measured gene expression trend in Mild (left half-arch) and SC (right half-arch). (**B**) Heatmap showing compounds recurrently reverting gene expression signatures of severity across the cell lines tested in the CMap database. Only drugs resulting in a significant one-sided Fisher’s test false discovery rate (FDR < 0.05) in at least three cell families were selected. The colour code represents the enrichment test FDR.

We then focused our attention on drugs able to interfere specifically with the RAGE pathway. By interrogating again the CMap database we identified two compounds with a significant impact on RAGE pathway in multiple cell lines, and significantly enriched amongst the top-scoring drugs for both severe and mild patients’ lists of genes: Anandamide and BRD-A15079084, an endocannabinoid derived from arachidonic acid and Phorbol 12-myristate 13-acetate, respectively. Post-treatment expression z-scores of both compounds showed an independent modulation of most of the RAGE receptor binding interactors, leading to an overall repression of their expression (Fig. S8A). We could not test the transcriptional impact of Azeliragon on this widest set of RAGE pathway genes, since it is not part of the compounds screened in the CMap database. Finally, we performed an unbiased analysis of the CMap database, seeking compounds potentially capable of reverting severity signatures across cell families (Fig 5B). This analysis yielded six hit compounds (NVP-BEZ235, SA-1973507, Simvastatin, Sotrastaurin, BRD-K69894866 and U-18666A) significantly recurrently predicted to rescue the transcriptional program underlying disease severity in at least three cell families (Fig. 5B). The reliability of this result was again tested via CMap drug labels’ reshuffling, showing that the likelihood of these drugs to be retrieved as significant hits by random chance is <10^-4^.

## Discussion

The molecular mechanisms that alter innate and adaptive responses leading to hyperinflammation, eventually resulting in acute respiratory distress syndrome (ARDS) and multiorgan failure in COVID-19 patients, remain to be fully elucidated. Our longitudinal single-cell multi-omic investigation uncovered a complex dynamics of COVID-19 immune response across time and disease severity. At admission we observed, independent of severity, a significant increase in IFN monocytes, a subset distinguished by the expression of IFN-stimulated genes (ISG) consistent with what was recently reported (*42*). At discharge, confirming previous observations (*19*, *43*), patients displayed a higher level of proliferating cytotoxic CD8^+^ T cells as compared to admission, reflecting the activation of adaptive immune responses over time. While this accrued across severity, the number of proliferating cytotoxic CD8^+^ T cells was higher in mild than in severe/critical COVID-19 patients already at admission, suggesting either a delay in the induction of CD8^+^ T proliferation or an active inhibition of their activity in more severe patients. Consistently, mild patients displayed, *vis a vis* severe/critical ones, higher levels of effector CD8^+^ T cells throughout hospitalisation and at one month after discharge. As to B plasma cells, their highest abundance was found at discharge in all patients, with a drop one month later. Notably, at discharge 100% of patients had measurable anti-SARS-CoV-2 IgG, *vis a vis* a more heterogeneous distribution at admission, confirming previous findings (*44*). While the presence of plasma cells in our cohort is consistent with the recovery of all enrolled patients, we probed whether different modalities or kinetics of immune response activation reflected disease severity and time to recovery. We thus investigated differential expression across cell families comparing disease severity across time, with a focus on distinct functional pathways increasing or decreasing linearly over time that yielded the following readouts: i) specific metabolic, mitochondrial and ribosomal signatures in T cells; ii) a significant enrichment for genes induced in response to type I interferon and genes implicated in the regulation of viral replication in IFN-monocytes from both mild and severe/critical patients at admission; iii) a strong increase over time, in severe/critical patients, of antigen processing and presentation related genes, consistent with a delayed rise of the adaptive immune response in these patients. Finally, the functional profiling of myeloid cells in our cohort revealed the RAGE receptor binding pathway as significantly enriched at admission in non-mild patients (p = 0,0142) with a general decreasing trend over time (p = 0.0282 for mild patients, p = 0.0073 for the others). RAGE belongs to the immunoglobulin superfamily and is expressed in several cell types including alveolar, neuronal, endothelial and immune cells (*45*–*48*). Of note, RAGE expression is increased in most of the pathological conditions associated with COVID-19 severity, such as ageing (*49*, *50*), diabetes (*51*), obesity (*52*–*54*), atherosclerosis (*55*), cancer (*56*), chronic obstructive pulmonary disease (*57*) and ARDS (*58*).

Myeloid cells are known to be permissive to SARS-COV-2 entry (*59*), although they do not express ACE2. Very recently, it has been reported that a small percentage of circulating monocytes and lung macrophages are infected by antibody-opsonized SARS-CoV-2 through the CD16 and/or CD64 receptor binding (*24*). Remarkably, this interaction led to inflammasome-caspase1 activation and pyroptosis, possibly contributing to the inflammatory sequelae observed in severe patients. However, in our cohort 75% of severe/critical patients, who clinically experienced high inflammatory features at the hospital admission, had no detectable (anti-S) IgG antibody titers. Thus, alternative mechanisms must be at play in driving systemic inflammatory responses by a direct interaction with circulating monocytes.

22 N-glycosides (*60*) and 17 O-glycosides (*61*) have been identified in the SARS-CoV-2 Spike trimeric structure; extensive glycosylation has been also observed at the interface between the S-RBD and ACE2 interaction, thus pointing to a glycan-based mechanisms, well beyond shielding, evolved by the virus for receptor recognition and effective infection (*62*). Starting from such evidence, we hypothesised the involvement of RAGE in SARS-CoV-2 recognition by myeloid cells. Through a combination of computational, biochemical and cellular assays, our results demonstrate RAGE-dependent entry of SARS-CoV-2 into monocytes and uncover a robust signature of RAGE pathway activation which is replicated across all major cohorts profiled worldwide at comparable resolution. RAGE is a multi-ligand receptor recognizing several damage-associated molecular pattern molecules (DAMPs). Our findings identify a novel role for RAGE in COVID-19 as a viral receptor. This result confirms the crucial importance of glycosylation sites for viral transmission and pathogenesis (*14*, *63*), and prompts us to reconsider the role of RAGE in viral diseases and host-pathogen interactions. Interestingly, human RAGE is encoded by the *AGER* gene that presents multiple single-nucleotide polymorphisms. Different studies have reported a direct link between *AGER* genetic polymorphisms and the severity of different pathological conditions, including Non-alcoholic fatty liver disease (*64*), cancer (*65*), Alzheimer disease (*66*) and cardiovascular disorders (*67*). Further analysis on the association between RAGE genetic variants and COVID-19 severity will be valuable for mapping patient susceptibility to develop damaging inflammatory signs.

Finally, we leveraged the scale of the Connectivity Map to identify candidate compounds that could be repurposed for COVID-19 treatment based also on the pathogenic mechanisms uncovered through our longitudinal study design.

First, given the transcriptional dynamics underpinning disease course (with a peak at admission and the resolution at post 1 month), we used the tendency of gene expression (high at admission, medium at discharge and low at post 1 month, and vice versa) to identify drugs that mimicked this pattern and thereby enhanced antiviral response. Strikingly, the only CMap compound, among those in current use for COVID-19 treatment, that had a significant effect on the interrogated COVID-related transcriptional signatures (p-value < 0.05 for B cells, NK cells and Myeloid cells) was Baricitinib. Of note, this drug was found to impact the RAGE pathway inducing a general reduction of its expression. Besides providing an independent validation on the centrality of RAGE to COVID-19 transcriptional dynamics, this analysis of the Connectivity Map revealed that several IFN-dependent interactors of the RAGE pathway have a trend of downregulation concordant with the healing course, both in mild and in severe/critical patients. This is consistent with the fact that for proper viral clearance, these genes must be strongly induced during the early stages of SARS-CoV-2 infection. Their uncontrolled activation is linked however with detrimental hyper-inflammation and cytokine storms (*68*). This double edge is consistent with the progressive decline in their activation over time, coinciding with the healing of both mild and more severe patients. Interestingly though, this general trend does not hold for the interferon regulatory factor 4 (IRF4), whose expression over time decreases in mild but increases in severe patients, likely reflecting different kinetics of activation in the two patient groups. As Baricitinib enhances the IRF4 pathway in both mild and more critical patients, further studies are needed to determine the optimal timing protocol for the potential use of this drug in patients with varying degrees of severity. Besides Baricitinib, we mined CMap to identify further drugs that could modulate the RAGE pathway, independently of the interrogated COVID-related transcriptional signatures. This led us to identify anandamide and BRD-A15079084 (or PMA), which affect RAGE-associated genes predicted by Ingenuity Pathway Analysis (IPA) (*69*) to be altered in COVID-19 patients. As we inferred similar effects on RAGE pathway modulation by these drugs in comparison with Baricitinib, we hypothesised that the structures of these molecules could be used as a baseline for the designing of new RAGE modulating drugs.

Next, we also pursued an unbiased complementary analysis aimed at identifying repurposable compounds that could revert SARS-CoV-2-induced transcriptional alterations in all the identified cell families. Our results revealed six compounds with evidence of a healing pathway-modulatory potential: NVP-BEZ235, SA-1973507, Simvastatin, Sotrastaurin, BRD-K69894866, and U-18666A. Among these, both U18666A and NVP-BEZ235 have been already reported to interfere with human coronavirus entry and viral production (*70*). In addition, the use of statins, the class of compounds to which simvastatin belongs, has been associated with lower COVID-19 patient mortality (*71*) and Simvastatin can downregulate the expression of RAGE (*72*, *73*), providing a mechanistic rationale for the further exploration of statin treatments in terms of timing- and RAGE-based stratification.

While our own evidence of the activation of the RAGE pathway was gathered from a cohort of COVID-19 patients likely infected with the same SARS-CoV-2 variant, both its replication in 13 published cohorts spanning multiple waves and our in-silico data strongly suggest a similar behaviour for the currently dominant Omicron variant. Furthermore, we cannot exclude that the activation of the RAGE signalling observed *in vivo* may be partially due to also binding by other alarmins, such as the S100 proteins (*74*). Indeed, recent evidence showed a systemic upregulation mainly of S100A8/9 in severe patients (*43*) and it was recently proposed that tocilizumab, currently used in the COVID-19 therapeutic regimen, may exert some of its effects by blocking S100A8/9 expression in COVID-19 (*33*). While additional work is needed to precisely partition the direct (SARS-CoV-2) versus possibly indirect (alarmins) contribution to RAGE activation in severe patients, the effect of tocilizumab on S100A8/9 further supports the relevance of the RAGE pathway as a target for therapeutic strategies for severe COVID-19 disease. Moreover, our data suggested the presence of an additional sensing of the SARS-CoV-2 by the RAGE receptor in monocytes, perhaps involved in the extrapulmonary post-acute sequelae of COVID-19 (*75*). To our knowledge, this is the first report revealing that RAGE might act as a novel SARS-CoV-2 sensor. Together, our high resolution longitudinal analysis of COVID-19 course uncovers the RAGE-pathway (so far held only as theoretical possibility (*76*)) as a new crucial route underlying COVID-19 severity and amenable to therapeutic targeting.

## Materials and Methods

### Clinical profiling of the cohort

20 adult patients who were admitted to the infectious Diseases Unit (IDU) of the University Hospital of Padua, Italy, between April and May 2020, were enrolled in the study. All patients were diagnosed with COVID-19, confirmed by SARS-CoV-2 reverse transcription-polymerase chain reaction (RT-PCR) testing of a nasopharyngeal swab. According to WHO guidelines, all patients were classified into mild, moderate, severe, and critical cases based on symptoms, clinical examination, and chest imaging. Demographic, clinical, laboratory data were extracted from paper and electronic medical records using a standardised data collection form. Laboratory data included: complete blood count, ESR, CRP, coagulation profile, serum biochemical tests and lymphocyte subpopulations. Both Chest X rays and Ct scan were also performed in all patients.

### PBMC isolation

Peripheral blood from enrolled controls and COVID-19 inpatients was collected in EDTA tubes and stored at 4 °C prior to processing for Peripheral blood mononuclear cells (PBMC) isolation. PBMC were isolated by density-gradient sedimentation using Ficoll–Paque PLUS (GE Healthcare, Germany) according to the manufacturer’s protocol. Post-purification the isolated PBMC were cryopreserved in cell recovery media containing 10% DMSO (Gibco), supplemented with 90% heat inactivated HyClone™ Fetal Bovine Serum (FBS; GE Healthcare, Germany) and stored in liquid nitrogen, until analysis.

### Single-cell multi-omics experimental pipeline

For all transcriptomics experiments, we took advantage of the BD Rhapsody Express platform. After thawing, PBMC from the 3-time point from each patient were labelled using Single Cell Labelling with the BD Single-Cell Multiplexing Kit (BD Biosciences, #633781) and 52 different BD AbSeq Ab-Oligos reagents following the manufacturer’s protocol (BD Biosciences). Three different timepoints (adm, disch, pst1mo) from two patients were pooled, washed twice, and resuspended in cold BD Sample Buffer (BD Biosciences) to achieve approximately 60000 cells in 620 mL (10000 cells per sample). Single cells isolated using Single Cell Capture and cDNA Synthesis with the BD Rhapsody Express Single-Cell Analysis System following the manufacturer’s protocol (BD Biosciences). After priming the nanowell cartridges, pooled samples from two were loaded into BD Rhapsody cartridges. Cell Capture Beads (BD Biosciences) were prepared and then loaded onto the cartridge according to the manufacturer’s protocol. Cartridges were then washed, cells were lysed, and Cell Capture Beads were retrieved. After retrieval beads were split in two, and different libraries were prepared. Firstly, we generated whole transcriptome analysis (WTA) cDNA libraries according to manufacturer instructions. Briefly, after reverse transcription, performed with BD Rhapsody cDNA Kit, Sample Tag and AbSeq sequences were separated from Cell Capture Beads by denaturation and Sample Tag and AbSeq libraries were generated with two PCR, according to BD protocol. Target cDNA on Cell Capture Beads was subjected to Random Primer Extension (RPE). RPE products were then amplified by PCR. All the three libraries were finally indexed with one of the 16 indexes for Illumina sequencing.

For the other half of the beads, cDNA underwent targeted amplification using the Human Immune Response Panel primers and a VDJ CDR3 supplemental panel via PCR, according to BD protocol. PCR products were purified, and mRNA PCR products were separated from sample tag and AbSeq products with double-sided size selection using Agentcourt^®^ AMPure^®^ XP magnetic beads (Beckman Coulter, #A63880). mRNA and Sample Tag products were further amplified. and purified using Agentcourt^®^ AMPure^®^ XP magnetic beads. Quality and quantity of PCR products were determined by using an Agilent 4150 TapeStation with High Sensitivity D5000 ScreenTape (Agilent). Final libraries were indexed using one of the 16 indexes for Illumina sequencing. A different index was used for each cartridge Quality of final libraries was assessed by using Agilent 2200 TapeStation with High Sensitivity D5000 ScreenTape and quantified using a Qubit Fluorometer using the Qubit dsDNA HS Kit (ThermoFisher, #Q32854).

### Sequencing

Different library types were pooled at different ratios based on their targeted reads per cell and the nanomolarity of the library pools was confirmed using the Agilent Bioanalyzer 2100. The library pools were sequenced on the NovaSeq 6000 (Illumina) loading the instrument with the concentration of 440 pM. WTA libraries were sequenced to achieve a minimum of 50.000 paired-end reads per cell for gene expression libraries, 26.000 for Abseq libraries and 480 for Sample Tag libraries. For the Immune response and VDJ panels: libraries were sequenced to achieve a minimum of 8.000 paired-end reads for Immune Response libraries, 26.000 for Abseq libraries, 480 for Sample Tag libraries, 3.000 for TCR libraries and 3.000 for B-enriched libraries.

### Single-cell Analysis

Single cell data will be made publicly available upon acceptance of the paper. All the bioinformatic analyses, and the code to reproduce the figures are available in a GitHub repository that will be made publicly accessible to ensure complete reproducibility and help the reader to consult, understand and re-use our data and analytical pipelines.

### Alignment and quantification

The demultiplexing of the raw data was performed bcl2fastq v2.20.0.422. The reads obtained from the demultiplexing were used as the input for the BD Rhapsody WTA and targeted analysis pipelines hosted on the Seven Bridges Platform. In detail, Reads have been aligned to the human genome GRCh38 with release 29 of GENCODE annotation, then collapsed to unique molecular identifier (UMI) counts and labelled by cell barcodes and sample tags. The result is a large digital expression matrix with cell barcodes as rows and gene identities as columns.

### Quality control, normalisation, batch correction, embedding and clustering

After getting the count matrices, standardised pipelines for filtering (cells with less than 800 genes expressed and more than 30% of mitochondrial reads have been filtered out), normalisation, dimensionality reduction, clustering and annotation of cell population were used (*77*). In particular, UMAP dimensionality reduction as implemented in Scanpy (*78*) was applied. Clusters were identified by applying the Leiden algorithm from Scanpy, which is a community detection algorithm that has been optimised to identify communities that are guaranteed to be connected. This resulted in clusters of cells that are more coherent with the biological phenotype and more reliably identify cell populations. The resolution parameter value was optimised by surveying the stability of the resulting clusters. This resulted in the identification of 32 clusters. Three of these clusters were isolated and then further sub-clustered to allow the assignment of a well-defined identity. Cluster annotation and integration with external references

Cluster annotation in immune cell types was obtained by a combination of the following approaches: i) Scanpy’s rank_genes_groups to identify the most characterising genes per clusters; ii) visualisation of specific immune markers for each cell population, iii) projection onto external references through ingestion ([link](https://scanpy.readthedocs.io/en/latest/generated/scanpy.tl.ingest.html)) a function implemented in Scanpy (*78*), to integrate embeddings and annotations projecting on a PCA that has been fitted on the reference data. In particular the single cell PBMC dataset from Ref. (*42*), including single cell multi-omics readouts from COVID-19 patients and controls using a similar single cell sequencing approach, was used as a reference.

### Differential abundance testing

Abundances of the different cell families and cell subclusters were compared across patient severity and sample time-points. Counts were analysed with the edgeR package (*28*, *79*, *80*), using a Negative Binomial Generalised Linear Model (GLM) that is suited to model over-dispersed data in the presence of limited replication.

### Pseudo-bulk differential expression analysis

The sc-RNAseq counts of each cell family, patient and sample time-point were summed together to generate pseudo-bulk counts, with the aim of obtaining more robust expression values while keeping the relevant information from the experimental design and the single-cell resolution. Separately for the two selected categories of disease severity (mild vs severe/critical), the pseudo-bulk counts were then fitted with a generalised linear model using the EdgeR package, to identify those genes characterised by a well-defined decreasing or increasing trend of the expression over the sample time-points.

### Functional enrichment analysis

The differentially expressed genes (DEG) of the pseudo-bulk analysis outlined above showing an absolute fold-change (FC) >= 2 and a false discovery rate (FDR) <= 0.05 were selected for further enrichment analysis in each immune cell type. The selected DEGs were used as input for the ‘string_db$get_enrichment’ (*81*) using Gene Ontology (GO) terms with 3 to 500 elements as background. Enriched GO terms with FDR <= 0.001 were then selected and the corresponding genesets investigated. Among these sets, the GO:0050786 genelist was then expanded using the Cytoscape ‘stringApp’ (*82*) in order to identify among the nearest neighbours with confidence score > 0.7 the ones showing the highest absolute FC values in Myeloid cells. These shortlisted genes were further investigated using the IPA software, starting from this list we have expanded the genes based on the connections categorised in IPA as discovered only in the immune system. This expanded list was then connected considering the same sources. The derived network was then overimposed with the DEG identified for both mild and severe and critical patients. This data was used as a source for a Molecule Activity Predictor (MAP) and the identified networks used for the subsequent drug repurposing analysis.

### RAGE pathway enrichment analysis

The enrichment of the RAGE receptor binding Gene Ontology Term (GO:0050786) was computed with a Gene Set Enrichment Analysis (GSEA) of relevant DEGs list. The DE analysis has been done as described previously by fitting the Negative Binomial GLM of EdgeR to pseudo-bulk expression values, comparing mild vs non-mild patients, and the pattern over time for mild and non-mild patients separately. The GSEA has been done with the clusterProfiler library (*83*, *84*), using gene lists ranked by the FDR of the differential analysis and the sign of the logFC.

### Cell culture

THP1 cells (TIB-202 ™ ATCC) were cultured in complete medium (RPMI1640 Cat.BE12-702F/12-Lonza, supplemented with 10% Foetal Bovine Serum (FBS) (Cat.10270-106-Gibco), 1% HEPES (Cat.BE17-737E-Lonza), 1% Penicillin-Streptomycin (Cat. DE17-602E-Lonza) at 37°C 5%CO^2^ until confluence and split 1:10 every 2-3 days. Primary human monocytes were obtained with the Pan Monocyte Isolation kit (130-096-537, Miltenyibiotec) following the manufacturer’s instructions.

Azeliragon toxicity was tested through an apoptosis assay. 1 × 10^5^ THP-1 cells were seeded on a 24-well plate in their culture medium. After 2 h of starvation, Azeliragon (2 or 4 μM) was added to cells. 2 μM Staurosporin (Sigma) was used as a positive control. Cells were incubated for 24, 48 and 72 hours at 37°C 5% CO^2^. Supernatants and cells were collected and stained with Annexin V APC (BD Pharmingen Cat#550475) according to the manufacturer’s instructions. Labelled cells were detected at FACS CelestaSorp (BD). Data are expressed as the percentage of Annexin V APC positive events. The gating strategy and the relative analysis were performed with FlowJo software.

### Molecular Modelling and Dynamics

The model of the RAGE protein was derived by the x-ray crystal structure 3O3U (*85*), while the reference type (RT) variant and the omicron variant models of Covid-19 RBD were both derived by the x-ray crystal structure 6ZGG chain B (*86*) (omicron mutations were introduced by mutating the relevant residues on the RT structure). The starting docking models for both S1-RBD variants interacting with the RAGE receptor were obtained from the highest-scoring configuration produced by using the LZerD webserver (*37*). The models were then solvated with fully atomistic TIP3P water, and Cl^-^ and K^+^ ions at a concentration of ~0.15 M in order to mimic the physiological ionic strength. MD simulations were carried on using the Gromacs 2020 package (*87*) and the Amber14SB force field (*88*), following simulation protocols similar to those we used in our previous works (*89*). Specifically, after energy minimization, we performed 200 ps of Simulated Annealing to allow side chains to equilibrate. We then performed two short simulations lasting 100 ps first in the NVT ensamble, and then in the NPT ensamble both with positional restraint on the heavy atoms of the protein. Finally, we performed an equilibrium MD simulation under periodic boundary conditions at constant pressure for 100 ns. Analysis was performed after 25 ns of equilibration. During the equilibrium MD simulation, temperature (T) and pressure (P) were kept constant at 300 K and 1 atm, respectively, using the Berendsen thermostat and barostat. Fast smooth Particle-Mesh Ewald summation was used for long–range electrostatic interactions, with a cut off of 1.0 nm for the direct interactions. Each simulation was performed in five identical replicas in order to check the results consistency and reduce the risk of being trapped by entropic barriers, thus improving the sampling of the configuration space available. Interaction probability was measured as the fraction of time in which the heavy atoms of listed Covid19-S1-RBD (or RAGE) residues were in close contact (distance < 3Å) with the RAGE (or Covid19-S1-RBD) protein during the MD simulation, as previously done in Ref. (*90*). Only the important interactions (interaction probability >60%) are shown.

### Binding Free Energy Computations

To produce reliable predictions of the binding free energy, we use the PRODIGY web server following the procedure used in our previous work (*39*). Specifically, the binding free energies are calculated as ensemble averages over the configuration space explored by the five different replicas. To speed up the calculation, we clustered the configuration space sampled by the various MD trajectories after equilibration (i.e., the last 75 ns each of the five replicas) according to their root mean square deviation (RMSD), and calculate the binding energy using one representative for 60 bigger clusters, being the clustering distance 1.3 Å. The final result is then obtained as the weighted average of the free energy computed for each of these configurations using the number of elements in the cluster as weight.

### Western Blot

Total protein extract was obtained with FASP buffer supplemented with cOmplete™, EDTA-free Protease Inhibitor Cocktail (Sigma-Aldrich). 30 ug of Protein extracts were separated by 4-12% Bold NuPage (ThermoScientific) and transferred onto PVDF membranes (BioRad). After blocking with 3% albumin (Sigma-Aldrich) and primary antibody incubation RAGE (ab3611, Abcam), ACE2 (XXX), ADAM17 (ab2051, Abcam)), TMPRSS2 (ab109131, Abcam), the membranes were incubated with an anti-rabbit peroxidase-conjugated secondary antibody (GE healthcare). Chemiluminescence was obtained by the ICL Substrate (GE healthcare), and images were captured with an imaging iBright (Thermo Fisher Scientific).

### Co-IP

For immunoprecipitation, 10×106 human peripheral blood monocytes cells were treated with SARS-CoV-2 Spike protein (RBD, His Tag) (GenScript) 100 ng/ml for 2 hours. After the treatment, the protein extraction was done with IP buffer (50 mM Hepes pH 7,5 (Lonza), 150 mM NaCl (Sigma-Aldrich), 1 mM EGTA (Sigma-Aldrich), 1,5 mM MgCl2 (Sigma-Aldrich), 10 mM NaF (Sigma-Aldrich), 10 mM Na2P2O7 (Sigma-Aldrich), 1 mM Na3PO4 (Sigma-Aldrich), 1% TritonX (Fluka), 10% Glycerol (Fluka), supplemented with cOmplete™, EDTA-free Protease Inhibitor Cocktail (Sigma-Aldrich). 500 μg of protein lysate was incubated with Anti-6X His tag^®^ antibody [HIS.H8] (ab18184, Abcam) overnight at 4°C, anti-Mouse IgG (Invitrogen) was used as isotype control. The protein complex was precipitated with Pierce™ Protein G Agarose (Thermo Fisher Scientific) for 2 hours at 4°C. The immune complexes were analysed by Western blot analyses with Anti-RAGE (ab3611, Abcam), antibody.

### THP1 and Monocytes infection with SARS-CoV-2

THP1 cells were plated at 5×10^5^ cell/ml in 48-well plates in 200 μl of RPMI-1640 supplemented with 1% fetal bovine serum (FBS) (Euroclone). Twenty-four hours later, the drug Azeliragon (Aze) was added in a range of concentrations from 1, to 4μM; 10μM Angiotensin II (AngII) was added alone or in combination with 2 μM of Aze. After 30 min, 20 μl of SARS-CoV-2 (*14*) (kindly provided by prof. Nicasio Mancini, Vita-Salute University, Milan, Italy) were added to obtain three multiplicities of infection (MOI): 1, 0.1, 0.01. After 1 h of virus adsorption, 400 μl of RPMI supplemented with 10% fetal bovine serum was added. Cell culture supernatants were collected 24, 48, 72 and 144 h post-infection and stored at – 80°C until the determination of the viral titers by a plaque-forming assay in Vero cells. Primary human monocytes were plated at either 2×10^6^ or 6×10^6^ cells for RNA purification and TEM processing, respectively, in low-adhesion 6-well plates and incubated for 30 min with 2 μM of Aze immediately after purification. After 30 min, 60 μl of SARS-CoV-2 isolate (*14*) were added to obtain a MOI of 1. After 2 h of virus adsorption, monocytes were collected into 2.0 ml Eppendorf and pellet for 10 min at 1500 rpm. Pellets were washed once with PBS and either 1 ml of TRIzol was added for RNA samples or 1.2 ml of fixation buffer was added for TEM samples as described below.

### TEM

Samples were fixed with 2.5% glutaraldehyde in 0.1M sodium cacodylate buffer pH 7.4 ON at 4°C. The samples were postfixed with 1% osmium tetroxide plus potassium ferrocyanide 1% in 0.1M sodium cacodylate buffer for 1 hour at 4°. After three water washes, samples were dehydrated in a graded ethanol series and embedded in an epoxy resin (Sigma-Aldrich). Ultrathin sections (60-70 nm) were obtained with an Ultratome Leica Ultracut EM UC7 ultramicrotome, counterstained with uranyl acetate and lead citrate and viewed with a Tecnai G^2^ (FEI) transmission electron microscope operating at 100 kV. Images were captured with a Veleta (Olympus Soft Imaging System) digital camera.

### Plaque-forming assay

Vero cells were seeded at 5.0×10^5^ cell/ml in 24-well plates in 500 μl of 1 ml of Eagle’s Minimum Essential Medium (EMEM) supplemented with 1% FBS (complete medium). Twenty-four hours later, 10-fold serial dilutions of SARS-CoV-2 containing supernatants were added in 300 μl of complete medium. After 1 h of incubation, the viral inoculum was removed and methylcellulose (Sigma, 1 ml in EMEM supplemented with 5% FBS) was overlaid in each well. After 4 days of incubation, the cells were stained with 1% crystal violet (Sigma) in 70% methanol. The plaques were counted after examination with a stereoscopic microscope (SMZ-1500; Nikon Instruments) and the virus titer was calculated in terms of plaque-forming units (PFU)/ml.

### Immunofluorescence

500.000 cells/mL cells were seeded the day before in a complete medium in a 24 well non-tissue culture plate (Cat. 351147 Falcon). The following day, cells were pretreated or not with 2μM Azeliragon (Cat.S6415-Selleckchem) for 30 minutes before adding 100 ng/mL of Sars-CoV-2 spike protein (RBD, HisTag) (Cat. ZO3483-1-GenScript) or infected using Heat-inactivated SARS-CoV-2 (VR-1986HK, ATCC) at 4 TCID50/mL for 2h at 37°C 5%CO2. Cells were collected and centrifuged at 120g for 5 min with cytospin (MPW-223c, MPW) to prepare slides carrying 50.000 cells each. Slides were then fixed for 20 min. in Paraformaldehyde (PFA) 4% p/v (Cat.158127 Sigma-Aldrich). Cells were permeabilized in the permeabilization buffer (PBS with calcium and magnesium (Cat. P4417-100TAB-Sigma-Aldrich) plus 1% Bovine Serum Albumin (BSA) (Cat.A9647-500G-Sigma-Aldrich) and 0,02% NP-40 alternative (Cat.492016-100ML) for 1h at room temperature prior to overnight incubation at 4°C with primary antibody 1:100 (6xHisTag clone#HIS.H8 Cat.ab18184-Abcam or SARS-CoV-2 spike polyclonal antibody, GeneTex). Secondary antibody (AlexaFluor488 chicken anti-mouse IgG Cat.A21200-Invitrogen) was diluted 1:500 in PBS with calcium and magnesium and maintained for 1h room temperature. Nuclei were counterstained with Hoechst 33342 (Cat.H3570-Thermo Scientific) for 15 minutes at room temperature. Images were acquired with Zeiss LSM800 confocal microscope and analysed with FIJI software.

### Internalisation assay

Fresh primary human monocytes were pretreated or not with 2μM Azeliragon (Cat.S6415-Selleckchem) for 30 minutes before being exposed with 100 ng/mL of Sars-CoV-2 spike protein (RBD, HisTag) (Cat. ZO3483-1-GenScript) at different times (5, 15, 30 and 60 minutes). Cells were collected and stained using primary RAGE antibody 1:100 (PA5-24787, Thermo Scientific) for FACS analysis. Samples were read with BD FACSCelesta Flow Cytometer (BD Bioscience) and analysed with FlowJo V10.0 software.

### Drug repurposing

The CMap database (*91*) was interrogated through the Query tool on the clue.io web portal (QUERY [clue.io]). Up- and down-regulated lists of genes were given in input to obtain compounds reverting severity signatures for each cell family separately or inhibiting RAGE pathway. Where gene signatures comprised more than 150 genes, the 150 most strongly differentially expressed (up- and down-regulated) were kept (150 is the maximum length allowed for a query). Outputted connectivity score tables were downloaded from the web portal and processed to identify compounds recurrently displaying a positive connectivity score and FDR<0.01 across screened cell lines. A one-sided Fisher’s test was performed for each compound on each cell family separately, and only drugs with Fisher’s FDR<0.05 in at least 3 cell families were selected.

For the analysis of drugs in current clinical use, the CMap database was interrogated similarly with the list of genes changing expression longitudinally. The output was filtered for the three compounds Dexamethasone, Baricitinib and Ritonavir and processed to test whether these compounds recurrently display a positive connectivity score and FDR<0.01 across screened cell lines (one-sided Fisher test).

For the RAGE-pathway focused analysis, a similar approach was pursued using activated and deactivated genes obtained from IPA predictions. The z-scores indicating the effect of Anandamide and BRD-A15079084 on gene expression were obtained from the CMap Command tool (COMMAND [clue.io]), downloaded as a gdc file and averaged across all the cell lines with a significant drug-driven reversal of gene expression signature in both mild and severe patients. The z-scores indicating the effect of Baricitinib on gene expression were obtained from GSE70138 (files GSE70138_Broad_LINCS_Level4_ZSPCINF_mlr12k_n345976×12328.gctx and GSE70138_Broad_LINCS_inst_info.txt) and averaged across all the tested cell lines. The average z-score for RAGE-pathway genes decreasing expression longitudinally was compared with the average z-score of random genes with a one-sided Wilcoxon Rank Sum test.

Randomizations needed to compute empirical p-values quantifying hits significance were performed by reshuffling the CMap drug labels 10,000 times.

## Acknowledgments

We acknowledge DeBioImaging Facility, Dept. of Biology, Padua University and the Microscopy Facility of the Fondazione Istituto di Ricerca Pediatrica (IRP) Città della Speranza for support in image acquisition and analysis. We thank all the patients who participated in the study, the entire clinical staff of the Infectious Disease Unit of the Padova University Hospital.

## Author contributions

*In vitro* experiments and library preparation: RA, RRS, FM, NB, BM

Sequencing of transcriptomic libraries: PF, ER, LR

Computational analysis of transcriptomic data: MB, NC, FE, CZ, CEV

Docking and molecular dynamic simulation: DB, FZ

SARS-CoV-2 infection experiments: IP, SG, EV

Drug repurposing analyses: AS, FI

Patient enrolment and consent: CP, AC

Study supervision, data interpretation and manuscript writing: AV, GT

## Data and materials availability

The experimental data supporting the findings of this study will be made available upon acceptance of the paper. All the code used to produce the results of this study has been organised in a repository available at Github.

## Supplementary Materials

Supplementary Text

Figs. S1 to S8

Tables S1 to S7

References (92–101)

