## Supplementary materials for "RAGE engagement by SARS-CoV-2 enables monocyte infection and underlies COVID-19 severity"

### Longitudinal single-cell analysis reveals Spike-RAGE interaction in monocytes as a new route defining COVID-19 severity

R. Angioni, M. Bonfanti, N. Caporale, R. Sánchez-Rodríguez, F. Munari, A. Savino, D. Buratto, I. Pagani, N. Bertoldi, C. Zanon, P. Ferrari, E. Ricciardelli, C. Putaggio, S. Ghezzi, F. Elli, L. Rotta, F. Iorio, F. Zonta, A. Cattelan, E. Vicenzi, B. Molon, C.E. Villa, A. Viola and G. Testa

**This PDF file includes:**

Supplementary Text
Figs. S1 to S8
Tables S1 to S2

**Other Supplementary Materials for this manuscript include the following:**

Tables S3 to S7
MDAR Reproducibility

**Supplementary Text:**

**Analysis of RAGE pathway activation in monocytes from publicly available datasets**

To validate the RAGE pathway activation gene signature that is described in the main text, we repeated the analysis described in the Materials and Methods section ("RAGE pathway enrichment analysis") on several publicly available sc-RNAseq results.

To work on data in the same format and with the same annotation, we started from the datasets that have been identified and standardised by Tian et al. in Ref. (*20*). These authors thoroughly catalogued the public single-cell dataset from COVID-19 patients and processed the relevant ones, by aligning cell and patient metadata and standardising the file formats. The datasets that we considered in our analysis were originally obtained from Ref.s (*15*, *22*, *42*, *92*–*101*).

We downloaded the processed datasets from the portal https://atlas.fredhutch.org/fredhutch/covid/ and converted them from Seurat object format to AnnData, to work with Scanpy. We took the normalised sc-RNAseq count matrices, and followed this procedure:

1) selection of the cells from blood samples and annotated as myeloid (CD14 Mono / CD16 Mono / cDC1 and cDC2),

2) aggregation of the counts to compute pseudo-bulk expression values for each sample of the dataset (filtering samples for which the number of myeloid cells is less than 50),

3) DE analysis with EdgeR, comparing COVID-19 patients vs healthy controls, and a pattern of increasing severity using the mild, moderate, severe annotation of Ref. (*20*).

4) for each of the DEG lists, GSEA with the clusterProfiler library (*83*, *84*), using gene lists ranked by the FDR of the differential analysis and the sign of the logFC.

Fig. S4A shows the RAGE receptor binding gene score (computed as described in the text), for the relevant conditions and for each of the 13 datasets considered. The results of the enrichment test, reported in Fig. S4B, show a consistent enrichment of the RAGE receptor binding pathway for the patients and – among them – for the more severe forms of COVID-19.

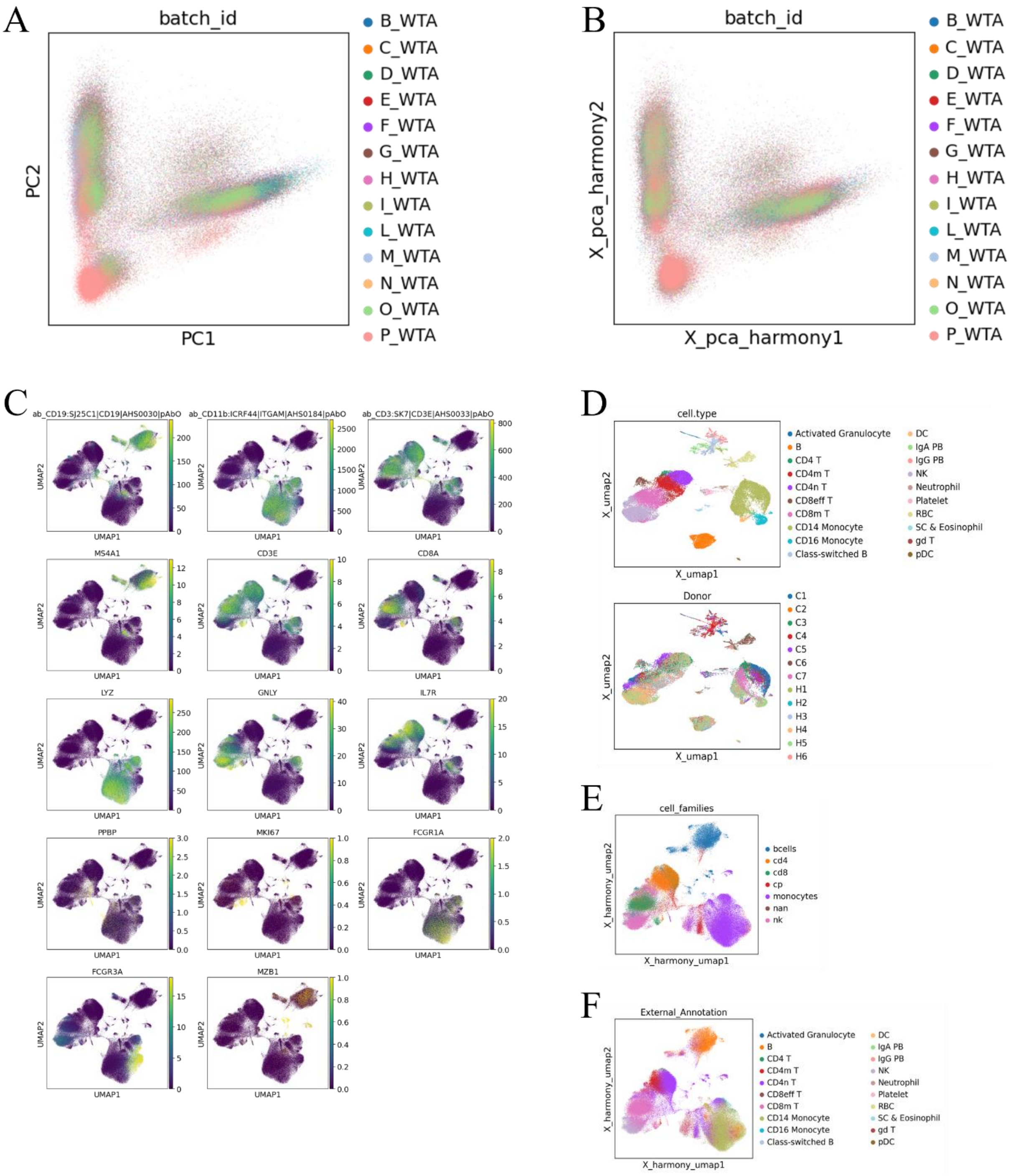


**Fig. S1. Integration and annotation of the COVID-19 patients single-cell transcriptomic data** (**A**) Overview of the batches for the single-cell transcriptomic experiment: cells from sc-RNAseq are shown in the reduced space of the first two principal components (PC1 and PC2). The cells are colored according to the experimental batch in which they were processed for library preparation and sequencing. Slight offsets between the single-cell distribution indicate the presence of a significant batch effect. (**B**) Batch-effect correction with Harmony: the same cells of panel A are now plotted in the space of the first two Harmony-corrected principal components (*X_pca_harmony1* and *X_pca_harmony2*). The distribution of the batches is now more interspersed because of the batch effect correction. (**C**) UMAP plots showing the quantification of a few surface proteins (CD19, CD11b and CD3) and the level of expression of several genes (MS4A1, CD3E, CD8A, LYZ, GNLY, IL7R, PPBP, MKI67, FCGR1A, FCGR3A, MZB1). These plots exemplify how both types of variables can be used to effectively identify cell populations of immunological relevance. (**D**) UMAP overviews of the cells of COVID-19 inpatients and controls from Wilk et al. (*42*). Cells are colored according to the cell type annotation (top panel) and the identity of the donor (bottom panel). (**E**) UMAP plot of the cell from our sc-RNAseq experiments colored according to the annotation of Wilk et al. projected to our dataset using the ingestion algorithm of Scanpy. Despite the differences between our experimental design and the one of Wilk et al., a substantial agreement is found between our cell type identification (see Fig. 1 of the main text) and the projected annotation. This finding confirms the robustness of our cell type annotation. (**F**) UMAP overview of our sc-RNAseq results showing some relevant cell annotations: sex of the patient, age interval of the patient (with a binning of 10 years), identity of the patient. The plots show a homogenous distribution of these covariates for the main clusters of the dataset.

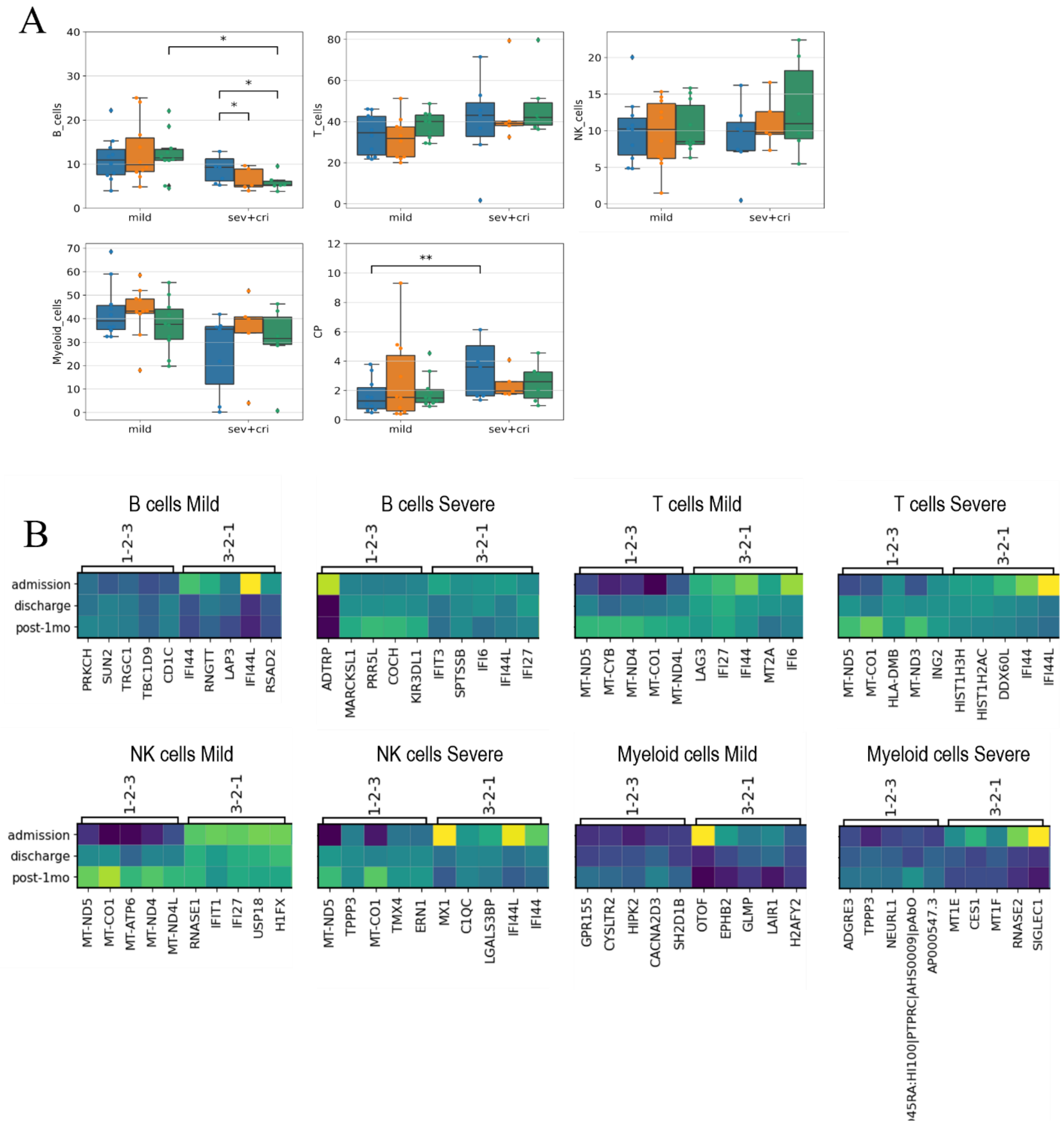


**Fig. S2. Differential analysis of the single-cell transcriptomic data.** (**A**) Box-and-whisker plots showing the significant changes of cell family abundance for the COVID-19 samples across time and patient severity. The box shows the dataset quartiles while the whiskers extend to the rest of the distribution, except for "outlier" points that are below/above the first/third quartile with a distance of more than 1.5 times the inter-quartile range. Differential abundance analysis was performed using the edgeR package (*28*, *79*, *80*), describing the data with the Negative Binomial

Generalised Linear Model (GLM). *$p \leq 0.05$; **$p \leq 0.01$; ***$p \leq 0.00$. (**B**) Matrix plots showing the average expression values across time for the top DEGs of both the increasing and decreasing longitudinal pattern, grouped by cell family and by patient severity. These plots exemplify the effectiveness of the adopted pseudo-bulk differential analysis in identifying genes that show the selected longitudinal pattern.

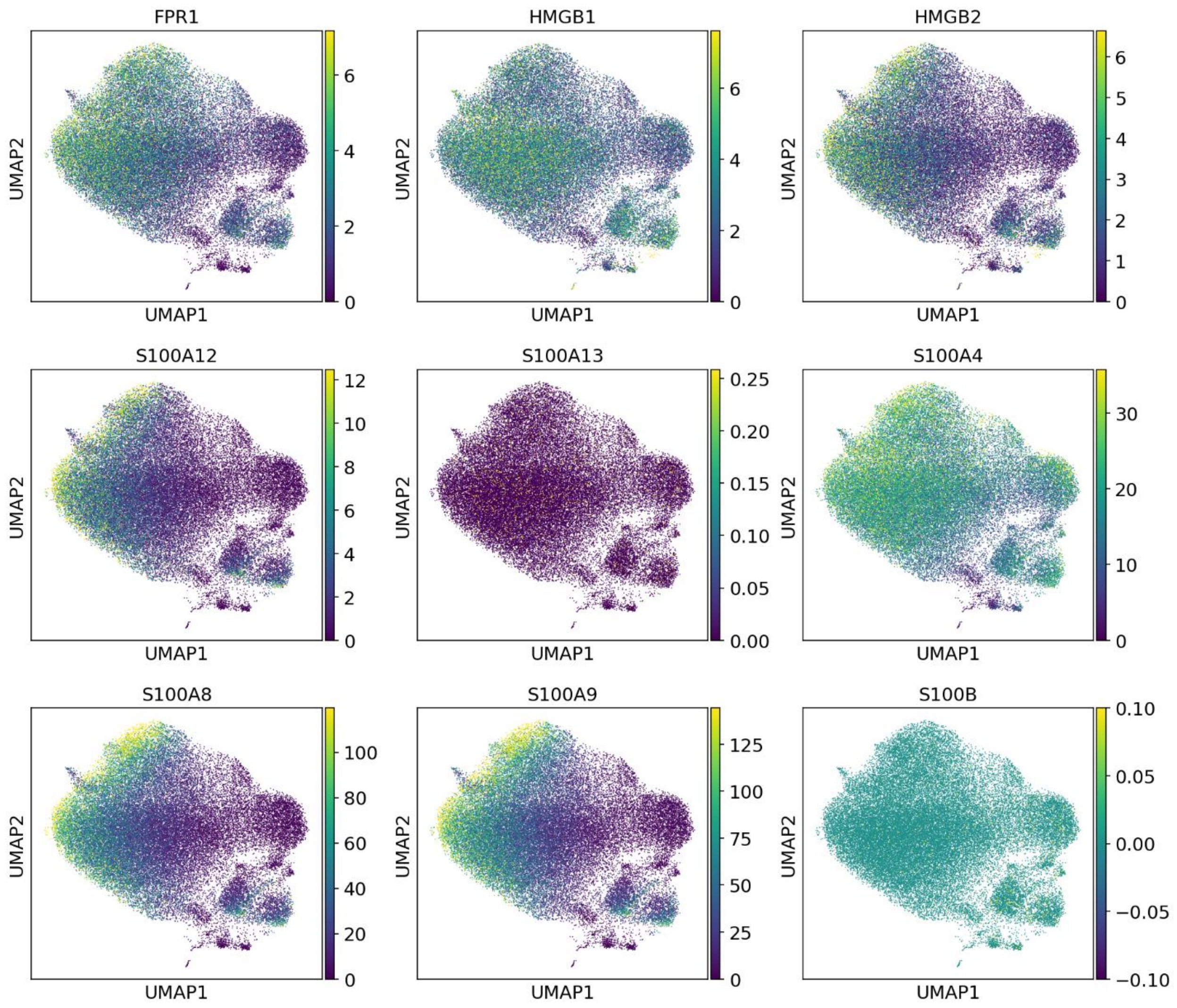


**Fig. S3. RAGE receptor binding genes.** UMAP plot of the single-cell expression of the genes included in the RAGE receptor binding Gene Ontology Term (GO:0050786). This gene list includes: FPR1, HMGB1, HMGB2, S100A12, S100A13, S100A4, S100A8, S100A9, S100B.

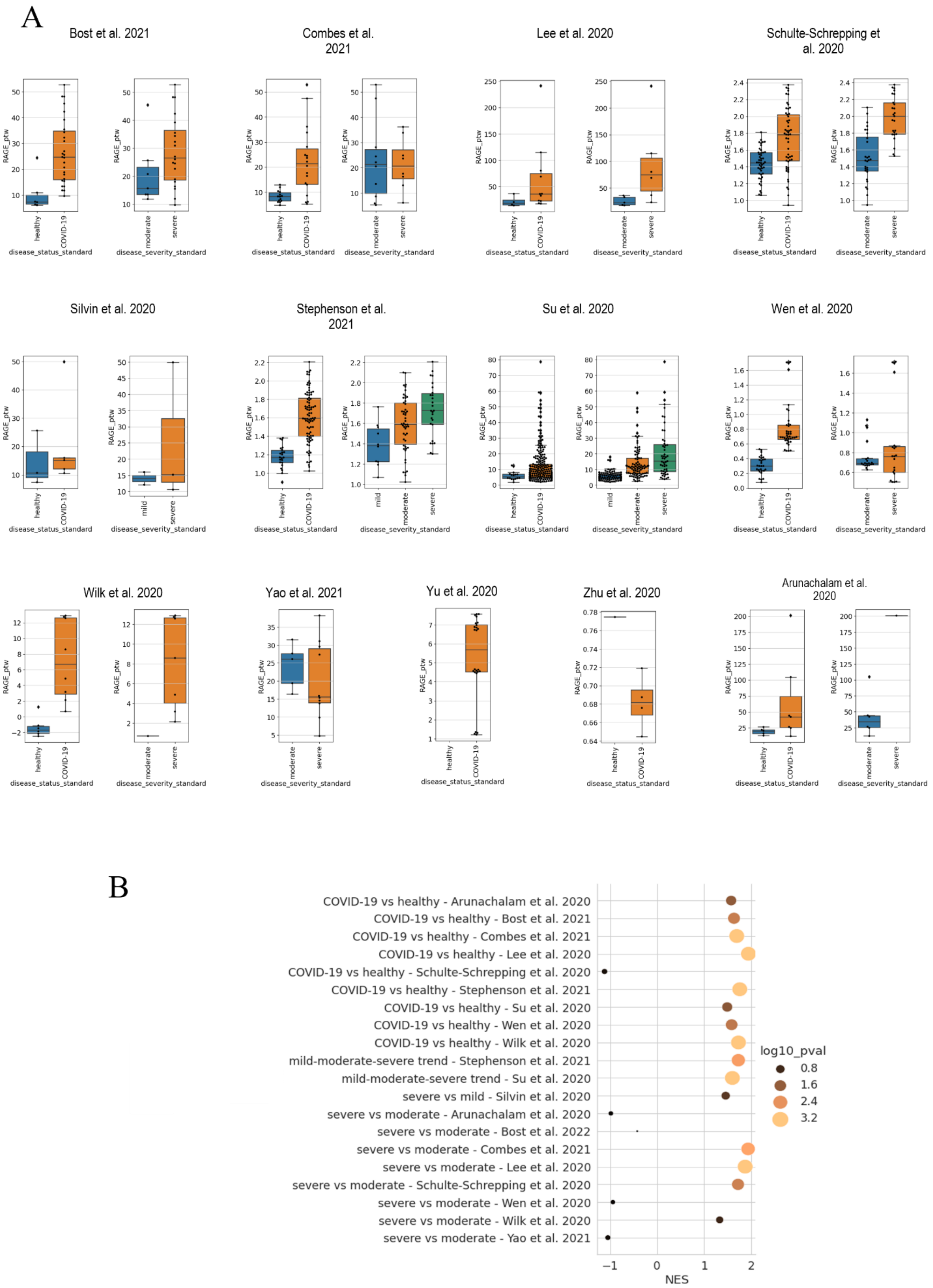
A
Bost et al. 2021
Combes et al. 2021
Lee et al. 2020
Schulte-Schrepping et al. 2020
Silvin et al. 2020
Stephenson et al. 2021
Su et al. 2020
Wen et al. 2020
Wilk et al. 2020
Yao et al. 2021
Yu et al. 2020
Zhu et al. 2020
Arunachalam et al. 2020
RAGE_ptw
healthy
COVID-19
mild
moderate
severe
disease_status_standard
disease_severity_standard
B
COVID-19 vs healthy - Arunachalam et al. 2020
COVID-19 vs healthy - Bost et al. 2021
COVID-19 vs healthy - Combes et al. 2021
COVID-19 vs healthy - Lee et al. 2020
COVID-19 vs healthy - Schulte-Schrepping et al. 2020
COVID-19 vs healthy - Stephenson et al. 2021
COVID-19 vs healthy - Su et al. 2020
COVID-19 vs healthy - Wen et al. 2020
COVID-19 vs healthy - Wilk et al. 2020
mild-moderate-severe trend - Stephenson et al. 2021
mild-moderate-severe trend - Su et al. 2020
severe vs mild - Silvin et al. 2020
severe vs moderate - Arunachalam et al. 2020
severe vs moderate - Bost et al. 2022
severe vs moderate - Combes et al. 2021
severe vs moderate - Lee et al. 2020
severe vs moderate - Schulte-Schrepping et al. 2020
severe vs moderate - Wen et al. 2020
severe vs moderate - Wilk et al. 2020
severe vs moderate - Yao et al. 2021
−1
0
1
2
NES
log10_pval
0.8
1.6
2.4
3.2

**Fig. S4. RAGE pathway enrichment in publicly available COVID-19 dataset.** (**A**) Comparison of the RAGE pathway gene score across condition (*COVID-19* vs *healthy*) and disease severity (*severe*: ICU patient, *moderate*: hospitalised patients not in ICU, *mild*: not hospitalised patients) for the myeloid cells extracted from 13 single cell studies previously integrated by Tian et al. in Ref. (*20*) (see Supplementary Text). The box-and-whisker plot shows the value of the RAGE pathway gene score for each of the samples averaged over the cells of the myeloid populations CD14 Mono, CD16 Mono, cDC1 and cDC2. The box and the whiskers are defined analogously to Fig 1B. (**B**) Results of the enrichment tests for the differential expression analysis performed on the COVID-19 public dataset. For each of the tests, the position of the dot on the x-axis indicates the value of normalised enriched score (NES) whereas the size and colour of the dot report the significance level of the test in terms of the negative logarithm of the p-value. NES and p-values have been computed as described in the Supplementary Text.

A

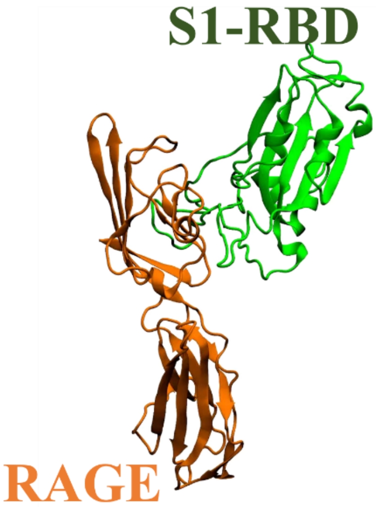


B

RAGE - RBD RefType

| Covid19-S1-RBD | | RAGE | |
|---|---|---|---|
| Covid19-S1-RBD amino acid | Interaction probability | RAGE amino acid | Interaction probability |
| TYR 449 | 99.3 | PRO 33 | 80.9 |
| ASN 450 | 75.5 | VAL 35 | 98.6 |
| LEU 452 | 91.7 | LYS 37 | 81.3 |
| THR 470 | 94.8 | SER 74 | 75.5 |
| GLU 471 | 65.6 | ARG 77 | 85.2 |
| ILE 472 | 61.9 | LEU 79 | 97.4 |
| VAL 483 | 67.2 | ASN 81 | 68.9 |
| GLU 484 | 87.1 | PHE 85 | 95.4 |
| PHE 490 | 99.9 | PRO 87 | 74.0 |
| SER 494 | 66.8 | | |

C

RAGE - RBD Omicron

| Covid19-S1-RBD | | RAGE | |
|---|---|---|---|
| Covid19-S1-RBD amino acid | Interaction probability | RAGE amino acid | Interaction probability |
| ARG 346 | 62.4 | ILE 26 | 84.2 |
| TYR 449 | 98.5 | LEU 34 | 82.1 |
| ASN 450 | 93.8 | VAL 35 | 99.3 |
| LEU 452 | 97.9 | SER 74 | 93.7 |
| THR 470 | 84.9 | ARG 77 | 95.9 |
| GLU 471 | 72.7 | LEU 79 | 86.0 |
| ILE 472 | 87.0 | PRO 80 | 66.0 |
| GLY 482 | 63.6 | ASN 81 | 80.3 |
| VAL 483 | 91.1 | PHE 85 | 99.1 |
| ALA 484 | 86.6 | PRO 88 | 93.0 |
| PHE 490 | 99.9 | | |
| SER 494 | 92.0 | | |

**Fig. S5. Interacting residues in the RAGE+Spike-RBD molecular dynamics trajectory** (**A**) Schematic representation of the interaction of the Spike-RBD and the RAGE receptor. Tables in (**B**) and (**C**) report the list of residues that take part in the interaction between the two proteins (RAGE and Spike-RBD reference type (B) or RAGE and Spike-RBD omicron variant (C)) and the percentage of the time in which the interaction is present along the 5 replicas of the MD trajectory. We consider a residue of one protein in interaction with the other protein when at least one of its heavy atoms is within 0.3 nm of a heavy atom of the other protein. We report only residues for which the interaction is stable along the dynamics (more of 60% of the time).

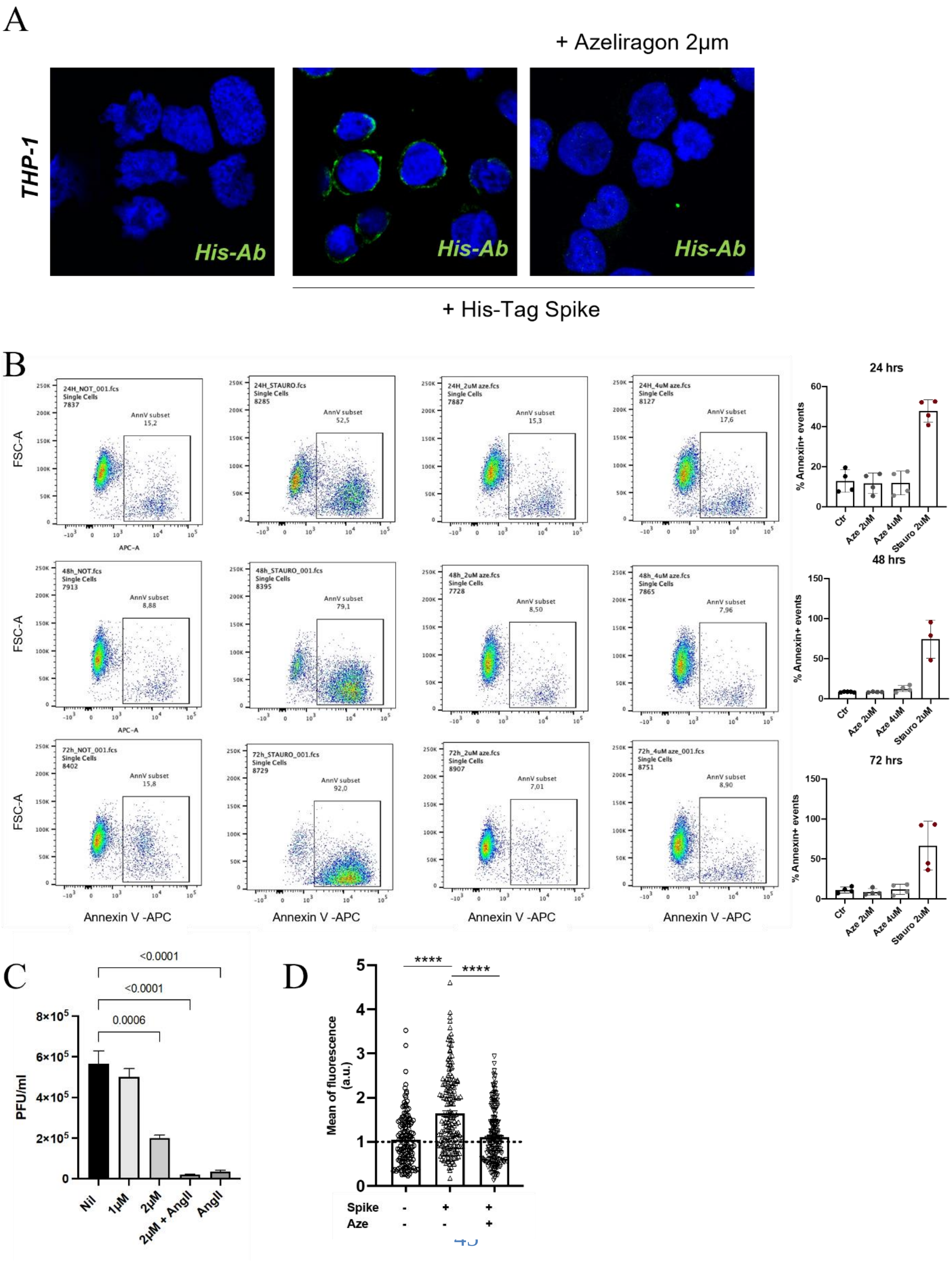
A
+ Azeliragon 2μm
THP-1
His-Ab
His-Ab
His-Ab
+ His-Tag Spike
B
24H_NOT_001.fcs Single Cells 7837
AnnV subset 15,2
24H_STAURO.fcs Single Cells 8285
AnnV subset 52,5
24H_2uM aze.fcs Single Cells 7887
AnnV subset 15,3
24H_4uM aze.fcs Single Cells 8127
AnnV subset 17,6
48h_NOT.fcs Single Cells 7913
AnnV subset 8,88
48h_STAURO_001.fcs Single Cells 8395
AnnV subset 79,1
48h_2uM aze.fcs Single Cells 7728
AnnV subset 8,50
48h_4uM aze.fcs Single Cells 7865
AnnV subset 7,96
72h_NOT_001.fcs Single Cells 8402
AnnV subset 15,8
72h_STAURO_001.fcs Single Cells 8729
AnnV subset 92,0
72h_2uM aze.fcs Single Cells 8907
AnnV subset 7,01
72h_4uM aze_001.fcs Single Cells 8751
AnnV subset 8,90
FSC-A
APC-A
Annexin V -APC
24 hrs
48 hrs
72 hrs
% Annexin+ events
Ctr
Aze 2uM
Aze 4uM
Stauro 2uM
C
<0.0001
<0.0001
0.0006
PFU/ml
Nil
1μM
2μM
2μM + AngII
AngII
D
****
****
Mean of fluorescence (a.u.)
Spike
Aze

**Fig. S6. Azeliragon effect on THP1 cells upon SARS-CoV-2 infection.** (**A**) Representative pictures of THP1 cells (Blue, nuclei were counterstained with Hoechst 33342) treated or not with 1-2μM Azeliragon (Aze) stimulated with His-Tag Spike (Green, Anti-6X His tag® antibody). (**B**) Flow-cytometry of THP-1 cells treated with Azeliragon (Aze) at growing doses for 24, 48 and 72 hours and stained with Annexin V APC (BD Pharmingen Cat#550475) according to the manufacturer's instructions. Staurosporin (stauro) was used as a positive control of apoptosis. Labelled cells were detected at FACS CelestaSorp. (**C**) Plaque forming unit (PFU) quantification after THP-1 cells infected with SARS-CoV-2 (0,01 MOI) for 72 hours in absences or presence of Azeliragon (1 and 2 μM) and Angiotensin II (AngII 10μM). (**D**) Quantification of the binding of His-Tag Spike to monocytes treated or not with 1-2μM Azeliragon (Aze) was measured as Mean of fluorescence, after 2 hours of stimulation. Data are presented as the mean of fluorescence intensity normalised on cells not infected. Nonparametric Mann-Whitney U test was used. $*p \leq 0.05$; $**p \leq 0.01$: $***p \leq 0.001$.

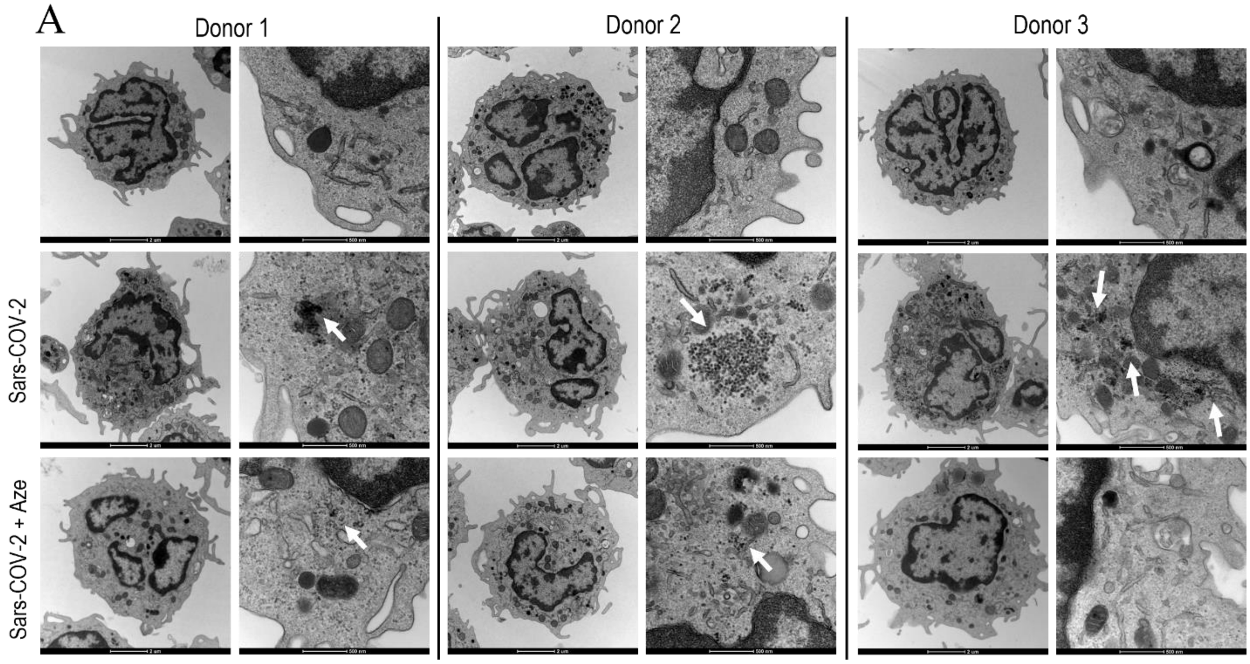


**Fig. S7. Representative TEM pictures of monocytes from the SARS-CoV-2 infection experiment.** Monocytes from 3 different donors exposed to replicative SARS-CoV-2 in the presence or absence of 2μM Azeliragon pre-treatment. The virions display a spherical shape (white arrow) and they are visible within the cytoplasm of infected monocytes.

A

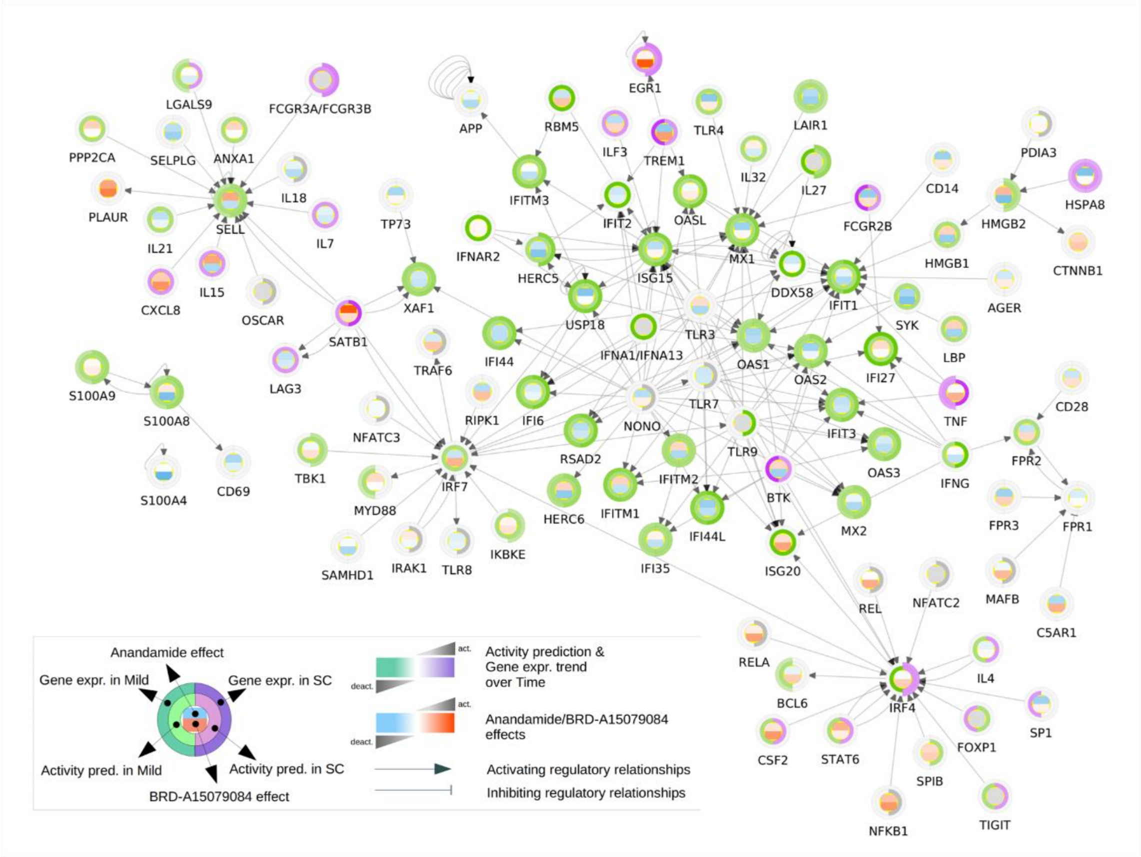


**Fig. S8. Extended network of the RAGE receptor binding interactors.** (**A**) In each node three metrics are reported: on the central heatmap the predicted effects of Anandamide (upper half) and BRD-A15079084 (lower half) on the gene expression; on the inner circle the predicted overall effect of interactors on the gene expression in Mild (left half-arch) and SC (right half-arch); on the outer circle the measured gene expression trend in Mild (left half-arch) and SC (right half-arch).

| | All patients n.20 |
|---|---|
| Age, years, median (IQR) | 56.5 (51.5-62.25) |
| Male, n (%) | 10 (50%) |
| Days from symptom onset to first positive test, days, median (IQR) | 4 (1-6.5) |
| Days from symptom onset and admission, days, median (IQR) | 6 (4-10.5) |
| Symptoms ad admission | |
| Fever, n (%) | 15 (75%) |
| Cough, n (%) | 12 (60%) |
| Severity of COVID-19 | |
| Mild, n (%) | 10 (50%) |
| Moderate, n (%) | 3 (15%) |
| Severe, n (%) | 4 (20%) |
| Critical, n (%) | 3 (15%) |
| C-Reactive Protein at the admission, mg/dl, mean (IQR) | 44.5 (2.75-57.75) |
| Comorbidities | |
| None, n (%) | 4 (20%) |
| 1-3, n (%) | 13 (65%) |
| >3 , n (%) | 3 (15%) |
| Anti-SARS-CoV-2 treatment | |
| Steroid, n (%) | 6 (30%) |
| Remdesivir, n (%) | 4 (20%) |
| Tocilizumab, n (%) | 0 (0%) |
| Chloroquine, n (%) | 11 (55%) |
| Admission to ICU, n (%) | 3 (15%) |
| Length of hospitalization, days, median (IQR) | 12 (5-15.75) |
| Patients discharged, n (%) | 19 (95%) |
| Death, n (%) | 1 (5%) |

**Table S1.** Demographic and Clinical data of COVID-19 patients at the hospitalisation admission.

| Agent | Use |
|---|---|
| Remdesivir | Antiviral; oxygen requirement and/or hih risk of worsening |
| Dexamethasone | Anti-inflammatory; oxygen requirement |
| Tocilizumab | Anti-IL-6 Ab; anti-inflammatory; rapidly progressing severe disease such as at initiation of aggressive supportive care |
| Baricitinib | JAK ½ inhibitor; anti-inflammatory and antiviral; rapidly progressing severe disease |
| Anakinra | Anti-IL-1 Ab; anti-inflammatory; hospitalised pts, with pulmonary involvement but with Pa/FiO2 >150 and suPAR >6 ng/ml |
| Sarilumab | Anti-IL-6 Ab; anti-inflammatory; hospitalized pts, in rapid clinical progression with need of mechanical ventilation |
| mAb: sotrovimab, carisivimab/imdevimab; etesevimab | Antiviral; nonhospitalized, not requiring oxygen, and at high risk for progression to severe COVID-19 disease |
| Molnupiravir | Antiviral; nonhospitalized, not requiring oxygen, and at high risk for progression to severe COVID-19 disease |
| Paxlovid (Nirmatrelvir/ritonavir) | Antiviral; nonhospitalized, not requiring oxygen, and at high risk for progression to severe COVID-19 disease |
| Remdesivir | Antiviral; nonhospitalized, not requiring oxygen, and at high risk for progression to severe COVID-19 disease |

**Table S2.** List of commonly used drugs in clinic for COVID-19 patients in Italy

**Table S3 (separate file).** Differential expression analysis of the sc-RNAseq data for the cells of each cell type cluster compared against all the other cells. Statistical analysis has been performed with Wilcoxon rank-sum test with Benjamini-Hochberg correction for multiple testing, as implemented in the Scanpy *rank_genes_groups* function (*78*). Only genes with corrected p-value $< 0.05$ are reported.

**Table S4 (separate file).** Differential expression analysis of the AbSeq data for the cells of each cell type cluster compared against all the other cells. Statistical analysis has been performed with Wilcoxon rank-sum test with Benjamini-Hochberg correction for multiple testing, as implemented in the Scanpy *rank_genes_groups* function (*78*).

**Table S5 (separate file).** Cell type abundance for each patient computed from sc-RNAseq data. Abundances corresponding to each patient are normalised to one.

**Table S6 (separate file).** Differential analysis of the sc-RNAseq data across the course of the disease for different patient severities. The DE analysis has been done by fitting the Negative Binomial GLM of EdgeR (*28*, *79*, *80*) to pseudo-bulk expression values, comparing the pattern over time for mild and severe+critical patients separately. FDR values were computed by correcting p-values for multiple testing using Benjamini-Hochberg method.

**Table S7 (separate file**). List of Gene Ontology terms with FDR $\leq 0.001$ in the functional enrichment analysis described in Materials and Methods. The table lists gene sets showing significantly increased or decreased expression trends over time in the mild and severe+critical patients.
